# Biosynthetic lanthanide-luminescent mini-proteins using genetic code expansion

**DOI:** 10.1101/2025.10.19.682977

**Authors:** Edan Habel, Haocheng Qianzhu, Elwy H. Abdelkader, Gottfried Otting, Thomas Huber

## Abstract

Non-canonical amino acids (ncAA) are promising as light-harvesting antennae for lanthanide luminescence in lanthanide-binding peptides and proteins. Here we present empirical insights into antenna-lanthanide interactions which reveal design principles of bright luminescent proteins. Peptides designed to act as lanthanide binding tags (LBT) show a trade-off between sensitization and lanthanide binding affinity. We generated a new protein, termed RF2, through computational design with nano-molar binding affinity and more than two-fold increase in terbium(III) luminescence. In this scaffold, 6-azatryptophan (6AW) achieved a ten-fold enhancement of the europium(III) luminescence *in vivo*. The RF2 6AW mutant also sensitizes the luminescence of dysprosium(III) and samarium(III). These results demonstrate the capability of *de novo* protein design to produce highly luminescent lanthanide-binding mini-proteins with a genetically encoded ncAA antenna.

## INTRODUCTION

Lanthanide ions are promising luminescent probes: they exhibit long luminescence lifetimes (0.5–3 ms), large pseudo-Stokes shifts (150–300 nm) and narrow emission bands.^1^ These photophysical properties are attractive for time-resolved fluorescence imaging as used with great benefit in immunoassays.^2-3^ In biological systems, however, free lanthanides readily precipitate with phosphates and their toxicity warrants strong chelators. Most critically, the molar extinction coefficients of lanthanide ions are extremely low, typically ranging between 0.1–10 M^-1^cm^-1^, which makes direct excitation inefficient. Fortunately, the lanthanide luminescence can be enhanced indirectly through nearby organic chromophores. This phenomenon, known as antenna effect, involves the photoexcitation of the organic chromophore followed by energy transfer to the lanthanide ion, overcoming the problem of low intrinsic absorptivity.

Choosing an optimal antenna is difficult because the energy transfer mechanism is complex. After excitation of the chromophore to the singlet state (*S*_1_), lanthanide(III) (Ln^3+^) excitation can occur via multiple pathways.^1^ The canonical pathway involves intersystem crossing (ISC) to the long-lived triplet state (*T*_1_), which in turn excites the Ln^3+^ ion.^1^ Sensitization therefore depends on the ISC efficiency as well as the *T*_1_-Ln^3+^ spectral overlap, rather than the more easily measurable absorbance and fluorescence properties of the antenna. In addition, energy transfer is also possible directly from the *S*_1_ state to the Ln^3+^ ion, although this mechanism is usually disadvantaged by its much shorter (nanosecond) lifetime.^1^ Nonetheless, direct *S*_1_ sensitization can be significant when there is strong spectral overlap, which has been explicitly modelled in protein-Ln^3+^ complexes.^4^ Crucially, both mechanisms rely on the antenna being in close spatial proximity to the Ln^3+^ ion and overlapping spectral properties. Additionally, Ln^3+^ emission is readily quenched by nearby water molecules, necessitating the exclusion of inner-sphere waters to achieve appreciable emission, even when other factors are favorable.^5^ Given these intertwined variables, predicting an antenna’s performance *a priori* is challenging, prompting the present empirical approach to identify effective sensitizers.

Peptidic lanthanide binding tags (LBTs), developed by Imperiali and co-workers,^6–12^ are biological lanthanide chelators with high specificity and affinity to lanthanide ions. They were specifically designed to include a tryptophan residue to act as the antenna for terbium(III) (Tb^3+^) excitation, with the backbone carbonyl oxygen of the tryptophan coordinating the lanthanide and the indole ring near the Tb^3+^ ion to efficiently photo-sensitize the Tb^3+^ luminescence. Additional carboxyl groups from acidic sidechains and carbonyls from amide-containing sidechains prevent the direct access of water to the lanthanide ion, enhancing the luminescence lifetimes and integrated emission intensities. Among the canonical amino acids, tryptophan (Trp) has the highest molar absorption coefficient (∼5500 M^-1^cm^-1^ at 280 nm) and provides the best antenna. Tyrosine (Tyr) in this position has also been established to sensitize the Tb^3+^ luminescence. Neither Trp nor Tyr, however, are effective antennae for any other lanthanide ion than Tb^3+^.

Over the last decade, there has been increasing interest in biosynthetic molecules to detect, isolate and utilize lanthanides. In 2019, lanmodulin, a protein from a lanthanide-utilizing bacterium, was identified to have picomolar affinity for lanthanide ions.^13^ Lanmodulin, which is structurally analogous to calmodulin, presents a lanthanide chelating environment similar to the LBTs with one residue coordinating the lanthanide ion by a backbone carbonyl oxygen. Featherston *et al*. reported that mutation of this amino acid to tryptophan sensitizes a Tb^3+^ ion for excitation at 280 nm.^14^ Further studies with luminescent lanthanide binding proteins and peptides have been reviewed by Pazos and co-workers.^15^ However, few attempts have been made to generate a new lanthanide binding peptides or proteins explicitly for optimal luminescence properties.

Substituting canonical amino acids by chemically synthesized antenna has been shown to allow for different lanthanide ions to be excited. Reynolds *et al*. used solid-phase peptide synthesis to substitute a tryptophan with a non-canonical amino acid (ncAA) containing either an acridone or carbostyril sidechain. This enhanced the excitation of europium(III) (Eu^3+^) luminescence and, with acridone, the luminosity of Tb^3+^.^9^ Other peptide-based lanthanide-luminescent probes were synthetically modified to include a fluorescent moiety,^16–19^ although this approach tends to increase the distance between the antenna and the lanthanide ion. All of these peptides require specific chemical synthesis and none can be produced *in vivo*.

Genetic code expansion (GCE) enables the site-specific installation of spectroscopically active ncAAs during protein biosynthesis in *E. coli* by using orthogonal aminoacyl-tRNA synthetase/tRNA pairs with amber stop suppression.^20^ We recently expanded the repertoire of genetically encoded tryptophan analogues, opening a modular route to incorporate antennae directly in recombinantly produced polypeptides.^21–24^ The present work systematically explores the performance of these ncAAs (Figure 1) as substitutes of Trp at the lanthanide-proximal site.

**Figure 1.**
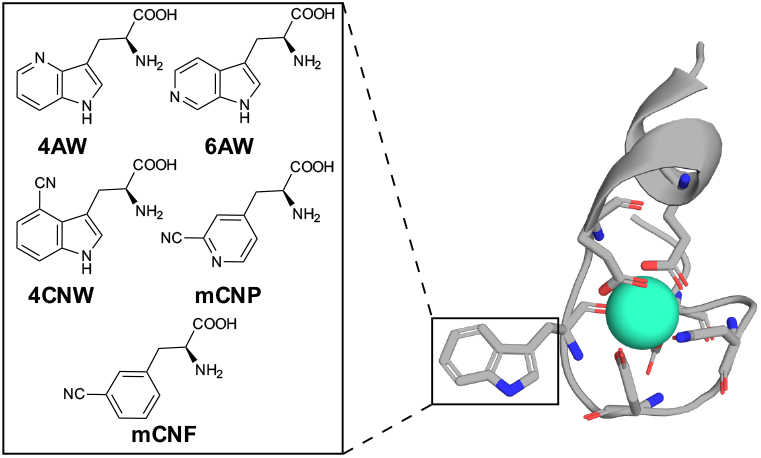
Binding geometry of the LBT (PDB ID: 1TJB) with a Tb^3+^ ion bound. The chemical structures are shown for the ncAAs used to substitute the antenna tryptophan, including 4-azatryptophan (4AW), 6-azatryptophan (6AW), 4-cyanotryptophan (4CNW), 3-cyano-4-pyridylalanine (mCNP), and 3-cyanophenylalanine (mCNF).

The ncAAs feature excitation wavelengths in the range 260–350 nm, which proved particularly beneficial for Eu^3+^ sensitization. We analyze the relationship between the antennae and the photo-properties of the Ln^3+^ complexes. Building on the experience gained with LBTs, we computationally designed three new lanthanide-binding mini-proteins. Two of these showed increased luminosity compared to their LBT parents and one displayed low-nanomolar binding affinity with Ln^3+^ ions. The best performing antenna was successfully combined with the most tightly binding lanthanide-binding protein, indicating that the luminescence enhancement achieved by the ncAAs is generalizable to other protein scaffolds, providing a basis for new *in vivo* probes.

## RESULTS AND DISCUSSION

All peptides and proteins were produced by recombinant protein expression in *E. coli*. ncAAs were installed using our previously developed two-plasmid system for *in vivo* incorporation of ncAAs via amber stop codon suppression (see Supplementary methods).^25^

### Enhanced luminescent properties with ncAAs

To obtain directly comparable results, we chose the previously reported lanthanide binding peptide with the sequence GFIDTNNDGWIEGDELLEG for its strong binding affinity and specificity for Tb^3+^, low intrinsic fluorescence and documented luminescence.^8^ The tryptophan at position 10 marks the antenna position, which was modified by ncAAs. The N-terminal glycine is a remnant of the N-terminal TEV cleavage site used to liberate the peptide following production as a fusion with the solubilizing NT* domain derived from spider silk protein.^25^

Following incorporation of different ncAAs into the LBT, we explored the link between their fluorescence and capacity for luminescence sensitization. Unexpectedly, the highly fluorescent 4CNW (fluorescence quantum yield >0.8 compared with ca. 0.15 in Trp)^26^ only minimally enhanced the luminescence of Eu^3+^ and delivered weaker integrated Tb^3+^ emission intensities. In contrast, the relatively weakly fluorescent 4AW revealed brighter luminescence intensity with Tb^3+^ than tryptophan and excited Eu^3+^. The best performance was achieved with 6AW, which features an increased fluorescence quantum yield.^24^ Compared with Trp, 6AW enhanced the Tb^3+^ luminescence 50% more and excited the Eu^3+^ luminescence more effectively than any of the other antennae studied.

Analyzing the lanthanide dissociation constant *K*_D_ of the LBTs, we found that the incorporation of the non-canonical antennae at position 10 in the LBT consistently reduced the binding affinity 2–8-fold (Table 1 and Figure 2). Given that the amino acid sequence of LBT was selected from a large peptide library using Tb^3+^ luminescence,^8^ it is unsurprising that even minor side chain modifications can reduce the ion affinity. In contrast, acridone and carbostyril sidechains installed in a similar LBT have been reported to perturb binding less (34 nM compared to 19 nM for the wild type).^9^ In our experiments, the *K*_D_ values differed up to 2.5-fold between Tb^3+^ and Eu^3+^, indicating extreme sensitivity of the lanthanide affinity towards optimal coordination geometry. While the nominal ionic radius differs by only 0.02 Å between Tb^3+^ and Eu^3+^,^27^ the effect may result from the non-isotropic electron densities of lanthanide ions.^28–29^

**Table 1.** Luminescence properties of the LBT-based Eu^3+^ and Tb^3+^ complexes^*a*^.

| Antenna | $\lambda_{\text{ex}}$<br>max<br>(nm) | Extinction<br>coefficient<br>at $\lambda_{\text{ex}}$ max<br>(M<sup>-1</sup> cm<sup>-1</sup>) | Quantum<br>Yield $\Phi_{S_1}$ | Intensity<br>Tb<sup>b</sup> | Intensity<br>Eu<sup>a</sup> | Apparent<br>$K_D$ (Tb<sup>3+</sup>)<sup>c</sup> (nM) | Apparent<br>$K_D$ (Eu<sup>3+</sup>)<sup>c</sup> (nM) | Lifetime<br>($\tau_{\text{Tb}}$)<sup>d</sup> | Lifetime<br>($\tau_{\text{Eu}}$)<sup>d</sup> |
| --- | --- | --- | --- | --- | --- | --- | --- | --- | --- |
| Trp | 280 | 5422 | 0.14<sup>e</sup> | 682 | 5 | 190 ± 40 | N.D. | 2.6 ms | 1.4 ms |
| mCNP | 267 | 1187 | – | 630 | 361 | 1300 ± 90 | 490 ± 30 | 2.7 ms | 1.4 ms |
| mCNF | 275 | 843 | – | 423 | 50 | 1400 ± 100 | 940 ± 100 | 2.7 ms | 1.4 ms |
| 4AW | 285 | 6272 | 0.01<sup>e</sup> | 728 | 86 | 1100 ± 50 | 580 ± 30 | 1.6 ms | 1.4 ms |
| 4CNW | 290 | 6041 | 0.80<sup>f</sup> | 323 | 5 | 1000 ± 40 | N.D. | 1.5 ms | 0.7 ms |
| 6AW | 260 | 2108 | 0.41<sup>e</sup> | 1072 | 378 | 800 ± 20 | 830 ± 20 | 2.7 ms | 1.4 ms |

<sup>a</sup> N.D. = not determined. <sup>b</sup> Integrated emission intensity for the complex with lanthanide ion measured at the excitation maximum and 545 nm for Tb<sup>3+</sup> and 615 nm for Eu<sup>3+</sup>. Samples contained 500 nM Ln<sup>3+</sup> ion with an excess of LBT in 20 mM MOPS pH 7.0, 50 mM KCl. Exact values are reported in Table S2. <sup>c</sup> Apparent $K_D$ values determined by titrating Ln<sup>3+</sup> (added as LnCl<sub>3</sub>) into a solution of 500 nM protein in 20 mM MOPS pH 7.0, 50 mM KCl. The $K_D$ values and error ranges were determined from a non-linear fit to the Hill equation for three technical replicates (see Supplementary methods). <sup>d</sup> Lifetime measurements were performed in 20 mM MOPS pH 7.0, 50 mM KCl at 500 nM Ln<sup>3+</sup> concentration. $\tau_{\text{Ln}}$ was determined by a non-linear fit of a single-exponential decay to lifetime measurements. The error for these measurements was < 0.01 ms. <sup>e</sup> From Abdelkader *et al.*,<sup>24</sup> measured in PBS. <sup>f</sup> From Qianzhu *et al.*,<sup>26</sup> measured in PBS.

**Figure 2.**
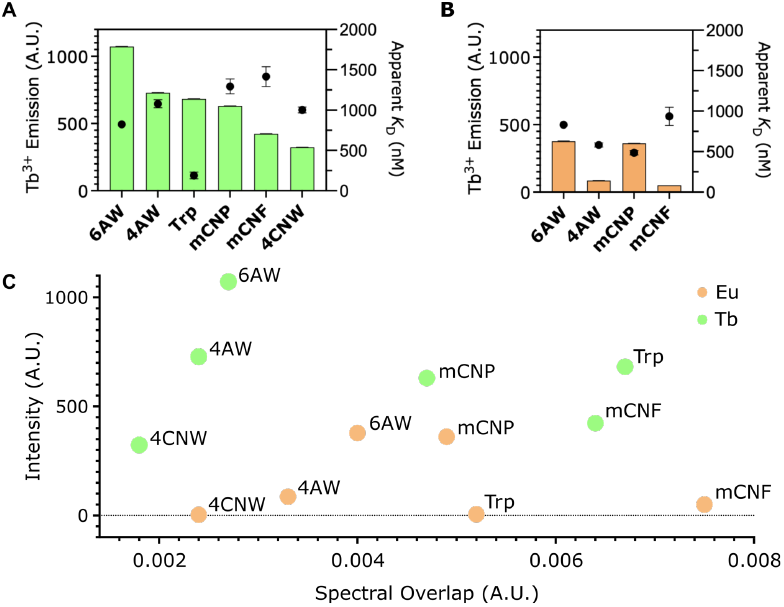
Luminescence properties of LBT variants. The histograms show the integrated emission intensities following excitation at the absorption maximum, which was 265 nm for 6AW, 285 nm for 4AW, 280 nm for Trp, 267 nm for mCNP, 275 nm for mCNF and 290 nm for 4CNW. The binding affinities for these complexes are indicated by black points which correlate to the right y-axis. (A) Tb^3+^ emission at 545 nm. The *K*_D_ value of the reference LBT is greater than reported,^6^ which we attribute to different buffer conditions. (B) Eu^3+^ emission at 615 nm. The signal intensity of LBT and LBT 4CNW was too weak to measure the affinity. (C) Plot of observed luminescence intensity versus spectral overlap for the series of LBT complexes. Spectral overlap was calculated as the integral of the scalar product between normalized absorption and emission spectra, with each spectrum area-normalized to 1.

Substituting the antenna with the less fluorescent mCNP greatly enhanced the Eu^3+^ luminescence with little altered enhancement of the Tb^3+^ emission. We speculated that the pyridine nitrogen of the mCNP amino acid directly coordinates the lanthanide ion, thus minimizing the interatomic distance between the antenna and the lanthanide ion. To test this hypothesis, we incorporated mCNF, which exhibits a fluorescence intensity similar to mCNP but lacks the pyridine moiety. mCNF still enhanced the luminescence of Eu^3+^ but significantly less than mCNP, suggesting that coordination plays a significant role. The dissociation constants in Table 1 support this interpretation. mCNP lends itself to rapid chemical diversification using the biocompatible nitrile-aminothiol arcNAT click reaction to generate a range of different fluorophores,^30,31^ which invites future exploration.

Our two most highly fluorescent antennae produced very different outcomes: 6AW was the most effective sensitizer for both Tb^3+^ and Eu^3+^ complexes, while 4CNW, despite its high intrinsic fluorescence, sensitized the luminescence of Tb^3+^ only modestly and hardly sensitized Eu^3+^. We also examined whether spectral overlap between the chromophore emission and Ln^3+^ absorption could predict the sensitization efficiency but found no simple correlation.

Coordination with hydration water efficiently quenches Ln^3+^ luminescence.^32^ As the luminescence lifetimes of the Eu^3+^ complexes were consistently 1.4 ms, the number of coordinated water molecules does not change between complexes with different ncAAs. The exception to this was LBT 4CNW, which showed a Eu^3+^ lifetime of 0.7 ms. In contrast, the Tb^3+^ luminescence lifetimes of the complexes with tryptophan analogues substituted at the 4-position (4CNW and 4AW) were significantly shorter (1.5 and 1.6 ms, respectively) than for the complex of the wild-type peptide or the LBT made with other ncAAs, suggesting a greater variation in the number of hydration water molecules coordinating the Tb^3+^ ion. Chemical changes at position 4 of the Trp antenna thus appear to alter the coordination geometry of the Ln^3+^ ion.

Table 1 show the absence of any simple relationship between the integrated emission intensity and the extinction coefficient or fluorescence quantum yield of the antenna. There was also no correlation with the spectral overlap between the antenna and the lanthanide ion. These findings underscore the complexity of the antenna effect in a peptide context and its dependance not only on chromophore brightness but also geometric and electronic compatibility with the lanthanide. The observation that the same antenna performed differently for Eu^3+^ and Tb^3+^ compromises any simple generalization across lanthanides. Furthermore, as all ncAAs reduced the binding affinity at least two-fold, we sought the design of a robust protein scaffold that tolerates antenna modifications without trade-offs in lanthanide binding affinity.

### Computational design of novel lanthanide-luminescent mini-proteins

To generate new, lanthanide-luminescent mini-proteins by computational design, we used Rosetta,^33^ RFDiffusion,^34^ and ProteinMPNN.^35^ The natural lanthanide-binding protein lanmodulin presents an attractive starting point as it features picomolar binding affinity for Ln^3+^ ions and has been studied in detail.^13,14, 36^ However, its four EF-hands, non-integer lanthanide binding stoichiometry of 3.5:1, and low expression levels in *E. coli* (<1 mg L^-1^) limits its suitability for a wider range of applications. We therefore used only the EF-hand 2 (EF2) of lanmodulin as the starting point of the protein design process. EF2 features a threonine residue with similar backbone carbonyl coordination to the lanthanide as the antenna Trp in LBT.^13^ To enable the coordination of the tryptophan carbonyl to the metal, the preceding residue is glycine.^7^ It is known that tryptophan substitutions at this position in lanmodulin enable lanthanide excitation without significantly perturbing the Tb^3+^ binding affinity.^14^

Coordinates of the 12 residues of EF2 (DPDKDGTLDAKE) were extracted from the NMR structure (PDB ID: 6MI5).^13^ Following mutating Thr7 to tryptophan, we energy minimized the side chain in Rosetta and used the resulting peptide geometry as seed structure input to RFDiffusion, diffusing a 70 or 50 amino acid protein with both N- and C-terminal extensions to the lanthanide ion binding motif. The generated backbone structures were used as input to ProteinMPNN, yielding 8 sequences for each backbone. A Tb^3+^ ion was placed for coordination in the protein and threaded with the ProteinMPNN sequences in Rosetta using the *ref2015* score function.^37^ The full details of the design process are reported in the supporting information.

Given the absence of explicit activity or binding considerations in RFDiffusion and ProteinMPNN, the scoring only accounted for a known binding geometry. Final structures were visually assessed, considering secondary structure diversity, solvent exposure of the Ln^3+^ ion, and plausible Ln^3+^-O distances. Three unique proteins with the best scores across all our evaluation criteria were chosen for experimental validation and expressed with an N-terminal His-tag and TEV cleavage site. In the following, these proteins are referred to as RF1, RF2, and RF3.

The three proteins have less than 35% sequence identity to lanmodulin. Using the Alphafold3 webserver,^38^ each sequence produced high-confidence structures with a calcium ion bound at the desired site. Each protein was recombinantly expressed in *E. coli*, using IPTG induction of the gene under a T7 promoter (see the Supplementary methods). RF1, RF2, and RF3 were isolated by immobilized metal ion chromatography (IMAC), with yields of 10 mg, 400 mg, and 15 mg per liter of cell culture, respectively. RF1 and RF3 not only expressed in much lower yields but were also more prone to precipitation at room temperature compared with the highly expressing RF2 protein.

All three designed proteins (RF1, RF2, RF3) produced a significant luminescence response when incubated with Tb^3+^(Table S5). Relative to the reference LBT, the luminescence intensities for RF1 and RF2 represented a 1.8-fold and 2.5-fold enhancement, respectively. In contrast, RF3 showed a diminished intensity of about 70 % compared to LBT. The Tb^3+^ affinities followed a similar trend, with RF2 showing the strongest binding affinity at *K*_D_ = 19 ± 3 nM, followed by RF1 at 320 ± 50 nM, and RF3 at 3300 ± 130 nM. RF1, RF2, and RF3 showed lifetimes of 1.39 ms, 1.45 ms, and 1.30 ms, respectively. All were above 1.2 ms, suggesting effective exclusion of water from the inner coordination sphere of the Tb^3+^ complex.^32^ However, the values are shorter than 2.5 ms, which is the typical lifetime of a fully shielded Tb^3+^ complex, implying water is not completely excluded. In summary, all data pointed to RF2 as the most attractive variant, and it was selected for further study.

Figure 3A and B shows the Alphafold3 model of RF2 with a Ca^2+^ ion (Figure 3A). The protein contains two α-helices and one β-sheet comprised of four β-strands. The EF-hand is positioned between β-strand 3 and α-helix 1. Trp42 is placed in the antenna position with the backbone carbonyl coordinated to the ion. A second aromatic amino acid, Tyr5, almost contacts Trp42 (Figure 3B). The presence of Tyr5 may be beneficial, as tyrosine excited at 280 nm can also excite lanthanide luminescence. In principle, the more distant Trp50 may also enhance the antenna effect. These two amino acids introduced by the sequence design of ProteinMPNN may help to produce a stable protein and we therefore retained them.

**Figure 3.**
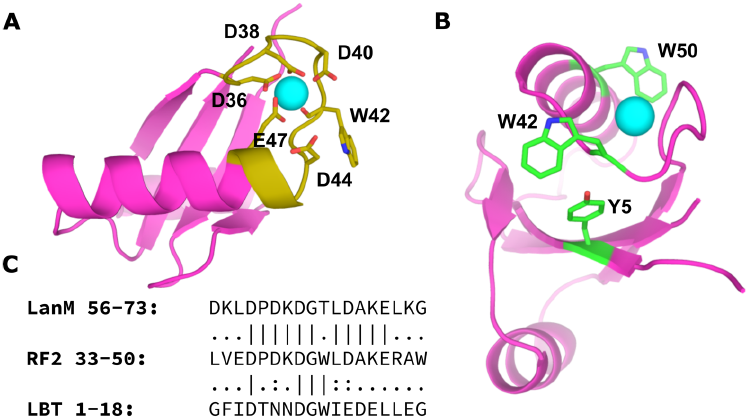
AlphaFold3 model of RF2. (A) RF2 model (magenta) with the EF-hand highlighted in gold, the Tb^3+^ ion in cyan, and the Tb^3+^ coordinating residues shown in stick representation. (B) Same as (A), but showing RF2 in a different orientation and showing the aromatic residues (green) instead of highlighting the EF hand. (C) Pairwise sequence alignment of lanmodulin EF2, RF2, and LBT with their flanking residues.

To confirm the modelled structure, we prepared a ^15^N/^13^C labeled sample of RF2 using ^15^N-labeled ammonium chloride and ^13^C-labeled glucose. The protein was isolated using IMAC with a yield of 100 mg purified protein per liter cell culture. Using 2D and 3D NMR experiments recorded on an 800 MHz spectrometer, we assigned the backbone chemical shifts (C, C_α_, C_β_, N, H, H_α_, H_β_) of most of the 71 amino acids in the protein complexed with yttrium (Y^3+^; Figures S12 and S13). Resonances of residues 10–14 could not be assigned. The backbone dihedral angles φ and ψ derived from the chemical shifts by the program TALOS+^39^ closely aligned with the Alphafold3 model (Figure S32).

### Enhanced brightness and more robust binding affinity with an ncAA antenna

We next tested if the increase in Eu^3+^ luminescence in LBT afforded by ncAAs is transferable to our designed protein. Replacing Trp42 with our best performing non-canonical amino acid, 6AW, we compared RF2 6AW with its wild-type parent RF2. The same two-plasmid system as used for LBT expression yielded 56 mg RF2 6AW per 1 L of cell culture.

Unlike LBT 6AW, RF2 6AW (λ_ex_ = 260 nm) showed no significant increase in Tb^3+^ luminescence relative to unmodified RF2 (λ_ex_ = 280 nm). We speculate that RF2 features an unusually high brightness owing to additional sensitization by Tyr5 and possibly Trp42, which are excited to a lesser extent at the absorption maximum of 6AW. In contrast, RF2 6AW showed a significant enhancement of Eu^3+^ luminescence, with the excitation spectrum showing increased excitation around 280 nm, relative to LBT 6AW (Figure S35). As the emission maxima for Tyr and Trp are at 304 nm and 350 nm, respectively, and 6AW possesses a second broad excitation band between 300 nm and 360 nm (Figures S11–S13), a non-radiative energy transfer from the nearby Tyr and Trp residues to 6AW can arguably add to the enhancement of the Eu^3+^ emission. A mutant with 6AW in position 50 and Trp retained at position 42 showed a decrease in Tb^3+^ excitation, and a minimal increase in Eu^3+^ excitation, suggesting that there is some sensitization from position 50 (Table 2).

**Table 2.**
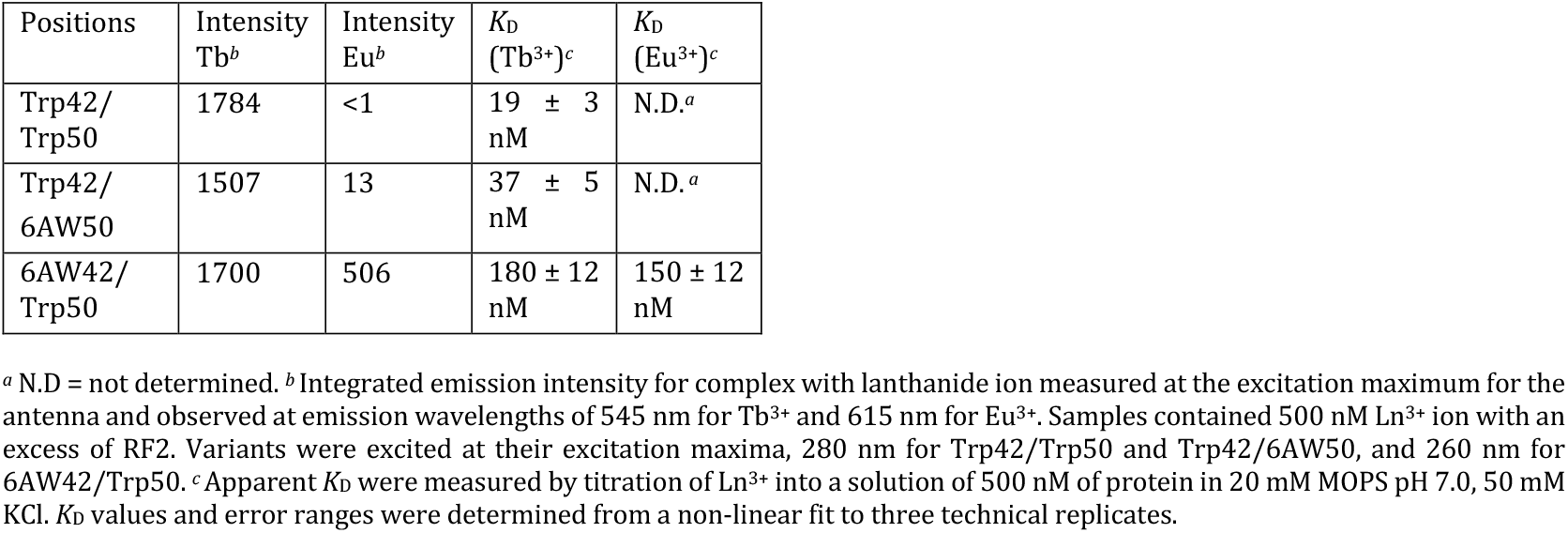
Luminescence properties of the RF2 Eu^3+^ and Tb^3+^ complexes^*a*^.

| Positions | Intensity<br>$\text{Tb}^b$ | Intensity<br>$\text{Eu}^b$ | $K_D$<br>($\text{Tb}^{3+}$)<sup>c</sup> | $K_D$<br>($\text{Eu}^{3+}$)<sup>c</sup> |
| --- | --- | --- | --- | --- |
| Trp42/<br>Trp50 | 1784 | <1 | $19 \pm 3$<br>nM | N.D.<sup>a</sup> |
| Trp42/<br>6AW50 | 1507 | 13 | $37 \pm 5$<br>nM | N.D.<sup>a</sup> |
| 6AW42/<br>Trp50 | 1700 | 506 | $180 \pm 12$<br>nM | $150 \pm 12$<br>nM |

<sup>a</sup> N.D. = not determined. <sup>b</sup> Integrated emission intensity for complex with lanthanide ion measured at the excitation maximum for the antenna and observed at emission wavelengths of 545 nm for $\text{Tb}^{3+}$ and 615 nm for $\text{Eu}^{3+}$ . Samples contained 500 nM $\text{Ln}^{3+}$ ion with an excess of RF2. Variants were excited at their excitation maxima, 280 nm for Trp42/Trp50 and Trp42/6AW50, and 260 nm for 6AW42/Trp50. <sup>c</sup> Apparent $K_D$ were measured by titration of $\text{Ln}^{3+}$ into a solution of 500 nM of protein in 20 mM MOPS pH 7.0, 50 mM KCl. $K_D$ values and error ranges were determined from a non-linear fit to three technical replicates.

Next, we measured the binding affinity of RF2 6AW with Tb^3+^ and Eu^3+^ and found that the ncAA affects the Ln^3+^ binding affinity by less than an order of magnitude (Table 2). 6AW at position 50 decreases the binding affinity only 2-fold, whereas 6AW at position 42 decreases the affinity almost 10-fold. As in the case of the LBT, the binding affinities differed between Eu^3+^ and Tb^3+^ (Table 2). Nonetheless, even the reduced binding affinities are in the nanomolar range, making the RF2 6AW construct suitable for a wide range of sensing applications.

Encouraged by the excellent enhancement of Eu^3+^ luminescence of RF2 6AW and its nanomolar binding affinities to Eu^3+^ and Tb^3+^, we recorded emission spectra with other trivalent lanthanide ions, including cerium, praseodymium, neodymium, samarium (Sm^3+^), gadolinium, dysprosium (Dy^3+^), erbium, thulium, and ytterbium. We identified weak, but significant, emission peaks with Sm^3+^ and Dy^3+^, which overlapped with the long-wavelength end of the emission peak from 6AW (Figure S26). While the Sm^3+^ emission was too weak for affinity titrations, we were able to determine the *K*_D_ value for Dy^3+^ (Figure S24) as 290 ± 30 nM, which corresponds to an only slightly weaker affinity than for Eu^3+^ or Tb^3+^.

To determine the number q of water molecules coordinated to the lanthanide ions, we performed luminescence lifetime measurements in D_2_O and used the modified Horrocks equation.^32^ Empirical correlation coefficients of luminescence quenching by H_2_O have been published for Tb^3+^ and Eu^3+^,^32^ as well as for Dy^3+^ and Sm^3+^.^40^ All four D_2_O titrations showed a clear linear response to the percentage of D_2_O (Figure S31), yielding q-values of 1.07, 1.06, 1.30, and 1.31 for Tb^3+^, Eu^3+^, Sm^3+^, and Dy^3+^, respectively. In view of the inherent uncertainty of this equation of ±0.3 for Tb^3+^ and Eu^3+^, and ±0.5 for Dy^3+^ and Sm^3+^,^40^ this indicates that the Ln^3+^ ion in the RF2 complex coordinates a single water molecule. With q-values close to 1 across Tb^3+^, Eu^3+^, Sm^3+^, and Dy^3+^, inner-sphere hydration (and thus OH-mediated quenching) varies only little between different lanthanide ions and ncAAs, allowing the brightness and apparent affinity trends to be attributed primarily to the antenna identity.

Two recent studies illustrate alternative strategies in lanthanide-luminescent protein design. A 2023 study grafted a Tb^3+^- binding EF-hand into a *de novo* immunoglobulin-like scaffold using ProteinMPNN, yielding a Tb^3+^ affinity of 8–12 µM by time-resolved luminescence.^41^ This establishes a *de novo* lanthanide-luminescent protein, but the moderate affinity limits its use under dilute or competitive conditions. More recent work focused on antenna modification, identifying acridone as a strong Eu^3+^ antenna if installed by GCE at three EF-hand positions of lanmodulin.^42^ Besides involving multiple antenna sites and four additional mutations, the construct leading to Ln^3+^ stoichiometries greater than 1:1, which compromises the analysis of antenna performance and lanthanide binding.

### New lanthanide-luminescent mini-protein shows remarkable selectivity and *in vivo* sensitization

Lanthanide ions have high charge density and bind promiscuously to anionic cellular components, including phospholipids,^43^ nucleic acids,^44^ metal-chelating proteins,^45^ and organic phosphates.^46^ Lanmodulin, despite its picomolar affinity to Ln^3+^ *in vitro*, is reported to remain largely unfolded and in its apo state in *E. coli* cells, even in the presence of 20 mM intracellular Y^3+^.^36^ The combination of low availability of free Ln^3+^ ions in cells with kinetically inhibited uptake in lanmodulin limits the utility of lanmodulin as an *in vivo* reporter.^47^ In addition, nonspecific binding of Ln^3+^ to DNA produces UV-excited luminescence,^48^ further confounding the luminescence readout. We therefore tested the RF2 scaffold for maintaining its luminescence signal in the face of competing cellular ligands and metal ions.

To investigate the utility of RF2 as a general intracellular lanthanide sensor, we measured the decrease in observable luminescence in the presence of a competitor,^49^ where the binding of various metal ions was evaluated by measuring the change in the sensitized luminescence of a solution containing the RF2/Tb^3+^ complex after the addition of an equimolar amount of competing metal ion. The remaining fraction of initial Tb^3+^ luminescence was used to determine the relative affinity of the competitor, with a fraction of ca. 0.5 indicating a binding affinity comparable to that of Tb^3+^, a fraction <0.5 indicating a stronger affinity, and a fraction >0.5 weaker binding.

The results for competing lanthanide ions were consistent with expectations based on previous titration experiments with RF2 6AW (Figure S25). The addition of Eu^3+^ resulted in a luminescence fraction of <0.5, while Dy^3+^ yielded a fraction >0.5 (Figure 4A). Along with many of the Ln^3+^ metals, we probed Ca^2+^ and Mg^2+^, which are hard cations like Ln^3+^ ions, and Zn^2+^ and Co^2+^, which typically prefer coordination environments with softer Lewis bases. None of the divalent metal ions decreased the Tb^3+^ luminescence, demonstrating their inability to compete for the Ln^3+^ binding site. Unexpectedly, however, Co^2+^ and Zn^2+^ increased the emission intensity (Figure S33), which may indicate the presence of a second metal binding site in RF2 that promotes the luminescence brightness by an allosteric effect. The strong preference for Ln^3+^ ions relative to the M^2+^ ions may be due to the difference in charge density or due to the subtle non-spherical nature of the Ln^3+^ ions fitting the designed pocket better.

**Figure 4.**
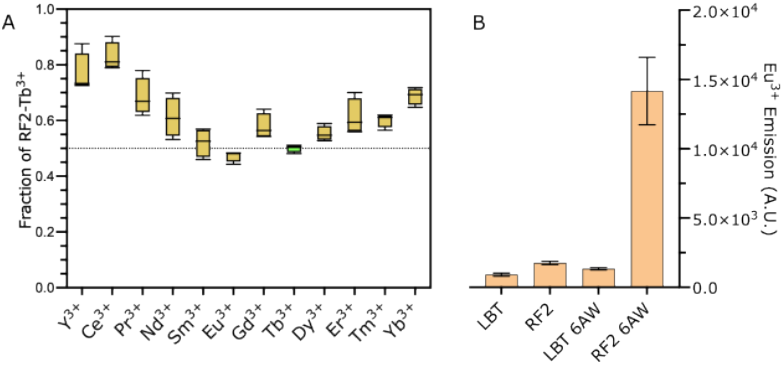
(A) Fold-change in the luminescence intensity (gold) of RF2 (1 µM) samples with Tb^3+^ (2 µM) and competitor (2 µM) in 20 mM MOPS pH 7.0, 50 mM KCl. Each data point represents the ratio of integrated emission intensity (collected at 545 nm emission with a 100 µs delay and 1 ms integration window, following excitation at 280 nm) normalized to the intensity of a sample containing only Tb^3+^. The green box indicates 50% of the intensity of the RF2/Tb^3+^ complex. Boxes indicate the mean and range of four technical replicates. (B) Eu^3+^ luminescence in *E. coli* B95 cells following overexpression of the protein constructs indicated. The emission values plotted were determined by integration of the emission intensity 600 µs after excitation at 280 nm and using a 1 ms integration window. Baseline luminescence was subtracted using cells not transformed with the respective plasmid(s) as blanks.

Interestingly, our competition assay gives rise to a parabolic trend in affinity across the lanthanide series, which is strongest for Eu^3+^ and Tb^3+^, with diminishing affinities towards both the heavier and lighter elements (Figure 4A). A similar curve has been reported for lanmodulin,^49^ and a minor deviation for Gd^3+^ has also been reported for LBT by Imperiali and coworkers.^7^ The improved sensitivity to lanthanide identity, relative to LBT, afforded by the RF2 mini-protein suggests the potential to design proteins with refined selectivity, which would constitute an important step toward biocompatible alternatives to current metal separation technologies.

In view of the selectivity of RF2 for Ln^3+^ over potential competitors found in biological matrices, we further tested its utility for detecting Ln^3+^ ions in the cytosol of *E. coli* cells. A culture of *E. coli* B95 cells was transformed with the dual-plasmid system. One plasmid encoded either the reference LBT, amber-LBT, or RF2 gene with an amber stop codon at either position 42 or 50. The second carried the orthogonal tRNA/tRNA synthetase pair for the incorporation of 6AW. The cells were grown to OD_600_ of 0.6 and protein expression was induced with 1 mM IPTG. After overnight expression, cells were resuspended in a solution of water containing 20 mM Ln^3+^, and then washed with 150 mM NaCl. The presence of the Tb^3+^- and Eu^3+^-complexes in the cells was readily detected (Figure S34). We chose to excite at 280 nm and 330 nm, which corresponds to the excitation maximum for tryptophan and the second excitation peak for 6AW, respectively. The luminescence of native cells with Eu^3+^ and Tb^3+^ excited at 330 nm featured a notably high background fluorescence due to the lanthanides binding to components in the cytosol that also sensitize their luminescence. Nonetheless, excitation at 280 nm produced strong *in vivo* sensitization in RF2 6AW, exceeding the signal of the other variants ten-fold (Figure 4B). This high signal-to-background enables single-wavelength, plate-based measurements in whole cells to detect the presence of Eu^3+^ in the cytosol.

## CONCLUSION

This study presents new methods for designing genetically encoded, bright, lanthanide-luminescent complexes in *E. coli* cells, leveraging GCE using selected aaRS:tRNA pairs. Our investigation of various antenna amino acids showed that an empirical approach is required for selecting good antenna for a range of lanthanides, because sensitization efficiency cannot be reliably predicted from the photophysical properties of the sensitizing moiety alone.

The computational designs yielded three proteins that all bind and excite Tb^3+^ ions, two of which outperform the reference LBT as a sensitizing complex. Our lead protein, RF2, has high expression levels, strong and specific binding to lanthanides, as well as enhanced luminosity. These peptides and proteins can be recombinantly produced and excited by a range of wavelengths (260–350 nm), and they retain their binding behaviors in the complex environment of the cellular cytosol. The genetically encoded RF2 6AW variant is, to our knowledge, the first report of a biogenic complex that sensitizes not only Tb^3+^ and Eu^3+^, but also Dy^3+^ and Sm^3+^ luminescence.

The characteristics of our novel, lanthanide-luminescent mini-protein extend the range of biological lanthanide applications. The design of Ln^3+^-binding proteins combined with the sensitization of Ln^3+^ by genetically encoded ncAAs opens the door to straightforward analysis of different Ln^3+^ ions in solution, including in intracellular environments.

## ASSOCIATED CONTENT

### Supporting Information

Experimental procedures for protein expression and purification; mass-spec of proteins produced; UV/Vis, fluorescence and NMR Spectra; equations for calculated values; titrations; analysis of secondary structures calculated from chemical shifts (PDF)

## Author Contributions

EH and HQ synthesized the protein constructs for this work. EH and GO contributed the NMR spectra and peak assignment. EH and EHA contributed the fluorescence and absorbance characterization of the amino acids. EH designed the sequences and contributed the characterization of the complexes. EH and TH wrote the original manuscript draft. GO and TH provided significant idea input into the writing of the manuscript. All authors contributed commentary or revised the final manuscript.

## ACKNOWLEDGMENT

The authors acknowledge the ANU Joint Mass Spectrometry Facility, the ANU Magnetic Resources Facility and the ANU Biopolymer facility. Financial support by the Australian Research Council for project funding (DP230100079, DP240100273 and CE200100012) is gratefully acknowledged.

## ABBREVIATIONS

aaRS: Aminoacyl-tRNA synthetase
GCE: Genetic code expansion
IMAC: Immobilized metal ion chromatography
IPTG: Isopropyl β-D-1-thiogalactopyranoside
ncAA: Non-canonical amino acid
WT: Wild-type

