## Supplementary Information for "Biosynthetic lanthanide-luminescent mini-proteins using genetic code expansion"

### Table of contents:

**Biosynthetic lanthanide-luminescent mini-proteins using genetic code expansion**

**Table of contents:**

**Supplementary methods**

- **a) General considerations**
- **b) Materials**
- **c) Expression and purification of LBT**
- **d) Intact protein mass spectrometry**
- **e) Luminescence measurements**
- **f) Integrated lifetime measurements**
- **g) Sequence design of EF2-derived mini-proteins**
- **h) q-value determination**
- **i) Lanthanide titration experiments**
- **j) Expression of isotope labeled RF2**
- **k) NMR measurements of $^{13}C/^{15}N$-labeled RF2**
- **l) Spectral overlap calculations**

**Supplementary figures**

- **Figure S1.** Intact protein mass spectrometry of LBT WT.
- **Figure S2.** Intact protein mass spectrometry of LBT 4AW.
- **Figure S3.** Intact protein mass spectrometry of LBT 6AW.
- **Figure S4.** Intact protein mass spectrometry of LBT 4CNW.
- **Figure S5.** Intact protein mass spectrometry of LBT mCNP.
- **Figure S6.** Intact protein mass spectrometry of LBT mCNF.
- **Figure S7.** Intact protein mass spectrometry of the EF2 variant RF1.
- **Figure S8.** Intact protein mass spectrometry of the EF2 variant RF2.
- **Figure S9.** Intact protein mass spectrometry of the EF2 variant RF3.
- **Figure S10.** Intact protein mass spectrometry of the RF2 42 6AW.

**Figure S11.** Intact protein mass spectrometry of the RF2 50 6AW.

**Figure S12.** HNCACB data of $^{15}N/^{13}C$-labeled RF2.

**Figure S13.** Overlay of $^{13}C$-decoupled 3D NOESY-$^{15}N$-HSQC and TOCSY-$^{15}N$-HSQC spectra of $^{15}N/^{13}C$-labeled RF2.

**Figure S14.** Absorbance spectra of the fluorescent ncAAs used in this study.

**Figure S15.** Spectral overlap of the Trp emission with the absorption of $Eu^{3+}$ and $Tb^{3+}$.

**Figure S16.** Spectral overlap of mCNP emission with the absorption of $Eu^{3+}$ and $Tb^{3+}$.

**Figure S17.** Spectral overlap of mCNF emission with the absorption of $Eu^{3+}$ and $Tb^{3+}$.

**Figure S18.** Spectral overlap of 4AW emission with the absorption of $Eu^{3+}$ and $Tb^{3+}$.

**Figure S19.** Spectral overlap of 4CNW emission with the absorption of $Eu^{3+}$ and $Tb^{3+}$.

**Figure S20.** Spectral overlap of 6AW emission with the absorption of $Eu^{3+}$ and $Tb^{3+}$.

**Figure S21.** Excitation scans of the LBT complexes with $Tb^{3+}$ and $Eu^{3+}$.

**Figure S22.** Binding affinity determination by titration of LBT variants with $Tb^{3+}$.

**Figure S23.** Binding affinity determination by titration of LBT variants with $Eu^{3+}$.

**Figure S24.** Binding affinity determination by titration of RF2 6AW complexes with $Tb^{3+}$, $Eu^{3+}$, and $Dy^{3+}$.

**Figure S25.** Excitation scans of the RF2 complexes with $Tb^{3+}$ and $Eu^{3+}$.

**Figure S26.** Fluorescence emission spectra of RF2 6AW with $Sm^{3+}$ and $Dy^{3+}$.

**Figure S27.** Luminescence lifetime measurement of RF2 with $Tb^{3+}$.

**Figure S28.** Luminescence lifetime measurement of RF2 6AW with $Eu^{3+}$.

**Figure S29.** Luminescence lifetime measurement of RF2 6AW with $Sm^{3+}$.

**Figure S30.** Luminescence lifetime measurement of RF2 6AW with $Dy^{3+}$.

**Figure S31.** $D_2O$ titrations of RF2/$Ln^{3+}$ complexes.

**Figure S32.** Comparison of the secondary structure of the AlphaFold3 model and the prediction made from the assigned chemical shifts.

**Figure S33.** Competition experiment with $M^{2+}$ ions.

**Figure S34.** *In vivo* sensitization of lanthanide luminescence.

**Figure S35.** Comparison of excitation spectra for RF2 6AW/$Eu^{3+}$, LBT 6AW/$Eu^{3+}$ and LBT/$Tb^{3+}$.

**Supplementary tables**

**Table S1.** Extinction coefficients of the amino acids used in this study

**Table S2.** Variables used in integrated lifetime calculations

**Table S3.** Amino acid sequences of the proteins used in this study

**Table S4.** Yields of different proteins co-expressed with G1(ncAA)RS in LB media with 1 mM ncAA

**Table S5.** EF2-RF variants with $Tb^{3+}$

**Table S6.** Determination of q-values from $D_2O$ titrations

**References**

### Supplementary methods

##### a) General considerations

No unexpected or unusual safety hazards were encountered. All buffers used for lanthanide luminescence measurements were treated with Chelex 100 resin. Metal stocks were made 10 mL volumetric flasks using water treated with Chelex 100 resin (BioRad), and a small amount of concentrated HCl to prevent hydroxide formation. Buffers used for experiments with lanthanides were also treated with Chelex 100 resin. Concentrations of rare earth element stocks were validated using complexometric titration with EDTA and Xylenol Orange (Sigma). For experiments with protein-$Ln^{3+}$ complexes, we used buffer conditions that have no or minimal affinity to lanthanide ions.[1]

##### b) Materials

The non-canonical amino acids 6-aza-tryptophan (6AW, Cat. No.: A861240) and 4-aza-tryptophan (4AW, Cat. No.: A264672), were purchased from Ambeed (USA). 4-cyano-tryptophan (4CNW, Cat. ID: V152071) was purchased from Advanced ChemBlocks Inc. (USA). 3-(2-cyano-4-pyridyl)alanine (mCNP, Cat. ID: CSMB00010457344) was purchased from Chemspace LLC (Kyiv, Ukraine). 3-cyano-phenylalanine (mCNF, Cat. ID: LC54307) was purchased from AK Scientific Inc., USA.

Buffer A: 50 mM Tris HCl pH 7.0, 300 mM NaCl
Buffer B: 50 mM Tris HCl pH 7.0, 20 mM imidazole, 300 mM NaCl
Buffer C: 50 mM Tris HCl pH 7.0, 500 mM imidazole, 300 mM NaCl
Buffer D: 10 mM Tris HCl pH 7.0, 50 mM KCl
Buffer E: 20 mM MOPS pH 7.0, 100 mM KCl.

##### c) Expression and purification of LBT

The gene for the fusion protein was ordered in a low-copy number pCDF plasmid under the control of a T7 promoter, either as the wild-type sequence (LBT-Trp) or with the tryptophan codon as an amber codon (LBT-Amb). The genes for the EF2 variants RF1, RF2, and RF3, as well as the RF2 amber mutants, were also cloned into the pCDF vector under control of the T7 promoter. High-copy number pRSF plasmids were used to encode the genes of the G1 tRNA synthetase mutants under the T7 promoter and the paired G1-tRNA under the *proK* promoter. This two-plasmid system has previously been established as an optimized system[2] to genetically encode ncAAs in *E. coli* B-95.ΔAΔ*fabR* cells.[3] For LBT-Amb, the two plasmids were transformed into cells concurrently. For LBT-Trp, the pRSF plasmid was omitted. Transformed cells were grown in LB media supplemented with 1 mM ncAA at 37 ˚C, and protein expression was induced with 1 mM IPTG at an $OD_{600}$ of 0.6. After 16–18 hours of expression, the fusion protein was isolated from the cells using immobilized metal-ion affinity chromatography (IMAC). This typically yielded 20–30 mg $L^{-1}$ of fusion protein. Incorporation of the ncAA was confirmed by mass spectrometry. The fusion protein was then concentrated to 1 mL in TEV buffer (50 mM Tris-HCl pH 8.0, 300 mM NaCl) and cleaved at 4 ˚C overnight after the addition of TEV protease at a 50:1 ratio of protein:TEV and 10 mM DTT. The resultant LBT-Amb fusion proteins were then isolated via size-exclusion chromatography (SEC) in volatile buffer (200 mM ammonium bicarbonate pH 8.5). Fractions containing the peptide were then isolated and lyophilized before luminescence analysis. For the EF2 proteins, IMAC elution fractions were dialyzed into TEV buffer and cleaved at RT overnight after the addition of TEV protease at a 1:1 ratio of protein:TEV and 2 mM β-mercaptoethanol. The EF2 proteins were then isolated by SEC in buffer D.

**d) Intact protein mass spectrometry**

Intact protein analysis was performed on an Orbitrap Fusion™ Tribrid™ mass spectrometer (Thermo Fisher Scientific, USA) connected to a Thermo Fisher Scientific UltiMate 3000 HPLC system equipped with ZORBAX 300SB-C3, 3.5 μm, 4.6 x 50 mm HPLC column (Agilent Technologies, USA). Approximately 50 pmol of sample was injected using a 500 μL/min linear gradient of solvent A (0.1% (v/v) formic acid in water) and solvent B (0.1% (v/v) formic acid in acetonitrile), ramping solvent B from 5% solvent B at the start to 80% after 12 min. Data were collected using an electrospray ionization (ESI) source in positive ion mode. Protein intact mass was determined by deconvolution using the program Xcalibur 3.0.63 (Thermo Fisher Scientific, USA).

**e) Luminescence measurements**

Luminescence measurements were performed on a Cary Eclipse fluorescence spectrophotometer (Agilent, USA) equipped with a phosphorescence module. Precise instrument settings for fluorescence measurements are indicated with the associated spectra. Time-resolved emission and excitation spectra were recorded with a 100 μs lag time, followed by a 1 ms integration time.

**f) Integrated lifetime measurements**

Luminescence lifetimes were measured on a Cary Eclipse fluorescence spectrophotometer (Agilent, USA) equipped with a lifetime measurement module. Each measurement employed a 100 μs delay followed by a 50 μs gate time, with both excitation and emission slit widths set to 20 nm. A 360–1100 nm filter was used to avoid signal from harmonic doubling. A total of 25 excitation flashes was used for a 15 ms decay window, and each dataset reflects the average of 50 repeated measurement cycles. Samples had a total volume of 600 μL in a 1 cm quartz cuvette. Samples were excited at their respective excitation maxima with emission recorded at 545 nm for $Tb^{3+}$ complexes and 615 nm for $Eu^{3+}$ complexes with a 20 nm slit width. Each sample was excited at the protein's excitation maximum with a 20 nm slit width and a recording of 500 nM $Ln^{3+}$ with no protein was subtracted. The instrument gain was set to 900. To account for variability in binding affinity, protein was iteratively added to the cuvette at small volumes (2 μL) until the initial intensity value did not change after addition of more protein. The lifetime measurement was then adjusted by subtracting the blank measurement, and then fit to the following exponential equation using GraphPad Prism, with the constraint on the fit that $K$>0.

$$Y = Plateau + (Y_0 - Plateau) \cdot \mathrm{e}^{-KX} \tag{1}$$

The values of $Y_0$, Plateau, and $K$ were then tabulated with a standard error of the fit. Residuals were inspected to assess the quality of fit. The integrated emission intensity was then calculated as:

$$I = \frac{Y_0 - Plateau}{K} \tag{2}$$

The standard error of the best-fit parameters was then propagated as follows:

$$\delta I = \sqrt{\left(\frac{1}{K}\right)^2 \cdot ((\delta Y_0)^2 + (\delta Plateau)^2\ ) + \left(\frac{Y_0 - Plateau}{K^2}\right)^2 \cdot (\delta K)^2} \tag{3}$$

Except for three very weakly luminescent samples, (RF2 6AW/$Dy^{3+}$, RF2/$Eu^{3+}$, LBT 4CNW/$Eu^{3+}$) propagated error did not exceed 1% of the integrated value. Exact values are tabulated in Table S2.

**g) Sequence design of EF2-derived mini-proteins**

Using PyMOL,[4] the coordinates of EF2 (residues 59–70) were extracted from the first frame of the lanmodulin NMR ensemble (PDB 6MI5).[5] Thr65 was mutated to tryptophan and the isolated segment was reindexed to begin at residue 1. The structure was energy minimized using the PyRosetta MinMover and the *ref2015* score function[6–7] The minimized coordinates were uploaded to the RFdiffusion Google Colab (v1.1.1).[8] Two sets of RFdiffusion backbones and sequences were generated using the built-in ProteinMPNN module. RF1 and RF2 were taken from set A with contigs set to "35:A1-15:24," and RF3 was taken from set B with contigs set to "20:A1-15:20." For both sets, iterations=50, num_designs=32, and symmetry="none" ProteinMPNN parameters were num_sequences=8, num_recycles=3, and mpnn_sampling_temp=0.1, with cysteine excluded from suggestions. Set A and B were downloaded and combined for processing. The final set included 64 sets of backbone coordinates, each with 8 associated sequences.

For each backbone, PyRosetta was used to mutate the coordinates to match each of the eight ProteinMPNN sequences. A $Tb^{3+}$ ion was placed 2.5 Å from the tryptophan backbone carbonyl along the C-O bond vector. Each model was minimized with *ref2015* and the final score was recorded as "rosettascore." The $C_\alpha$ alignment RMSD between lanmodulin EF2 residues 59–70 and the corresponding segment in each design was computed and recorded as "bb_RMSD." Distances between side-chain oxygen atoms and the antenna tryptophan carbonyl were measured, the count of O-$Tb^{3+}$ distances below 3.0 Å was recorded as "nO," and an RMSD to a 2.5 Å target for these O-$Tb^{3+}$ distances was calculated and recorded as "O_RMSD." The 2.5 Å target approximates an $Ln^{3+}$-O bond length in protein crystal structures containing lanthanides. The combination of rosettascore, bb_RMSD, nO, and O_RMSD was used to select one sequence per backbone. The resulting models were inspected in PyMOL and three designs were selected to represent different tertiary structures. For expression, the first residue was changed to glycine to introduce an N-terminal TEV site.

**h) q-value determination**

The q-value was determined using a version of the Horrock's equation, originally:

$$q = A\big((k_{H_2O} - k_{D_2O}) - B\big) - C \tag{4}$$

where A is a proportionality constant for the quenching of the lanthanide ion by O-H oscillators, *k* is the rate constant for the radiative decay of the lanthanide emission, parameter B is the corrective factor, accounting for other deactivating vibrations surrounding the ion, and parameter C is the outer-sphere contribution of closely diffusing solvent molecules.[9]

As the outer-sphere contributions are expected to be equivalent between the complex in $H_2O$ and $D_2O$, the difference between $k_{H_2O}$ and $k_{D_2O}$ is attributed primarily to the inner-sphere solvent. C was therefore omitted for all ions giving the equation:

$$q = A\big((k_{H_2O} - k_{D_2O}) - B\big) \tag{5}$$

For samarium, this equation is:[10]

$$q_{Sm} = 0.026(1/\tau_{H_2O} - 1/\tau_{D_2O}) \tag{6}$$

For europium, this equation is:[11]

$$q_{Eu} = 1.2((1/\tau_{H_2O} - 1/\tau_{D_2O}) - 0.25) \tag{7}$$

For terbium, this equation is:[11]

$$q_{Tb} = 5((1/\tau_{H_2O} - 1/\tau_{D_2O}) - 0.06) \tag{8}$$

For dysprosium, this equation is:[10]

$$q_{Dy} = 0.024(1/\tau_{H_2O} - 1/\tau_{D_2O}) \tag{9}$$

The lifetime in $D_2O$ was determined by plotting the lifetime of the $Ln^{3+}$-complex at mole fractions of $D_2O$ (0.2, 0.4, 0.6 and 0.8) followed by a linear fit to this data. The $\tau_{D_2O}$value was then determined by extrapolating to a $D_2O$ mole fraction of 1.0. These parameters are empirically determined based on the assumption that O-D oscillators effectively do not contribute to non-radiative deactivation, and that all other deactivation paths are the same between $D_2O$ and $H_2O$. The estimated uncertainties with using this technique are listed with the calculations of the q-values below.

**i) Lanthanide titration experiments**

Lanthanide binding affinity titrations were performed on a TECAN Infinite 200 Pro M Plex plate reader (Tecan, Switzerland) using black 384-well flat bottom plates (Corning, USA). Protein samples were prepared in Chelex-treated buffer E. Each well was a series of 1:2 serial dilutions of the corresponding lanthanide chloride stock, whose concentration was determined colorimetrically as described above. For each well, a background blank containing the same concentration of lanthanide ion was used for blank subtraction. Time-resolved luminescence measurements were performed for $Eu^{3+}$ and $Tb^{3+}$ with a 100 µs delay followed by a 1 ms integration window. For $Dy^{3+}$, time-resolved settings were not used, instead collecting the emission directly under steady-state conditions. Excitation wavelengths were matched to the known absorption maxima of each fluorinated amino acid residue, with emission measured at 545 nm for terbium, 615 nm for europium and 570 nm for dysprosium. Samples were equilibrated for 15 minutes at room temperature prior to measurement. Each point reflects the average of three replicate wells from replicate titrations. The averaged dataset was fit to the Hill binding equation using GraphPad Prism:

$$I = \frac{Bmax \cdot [Ln^{3+}]}{K_D + [Ln^{3+}]} \tag{10}$$

**j) Expression of isotope labeled RF2**

$^{15}N/^{13}C$-labeled RF2 was produced in a bioreactor (Infors-HT, Switzerland) at 300 mL media volume using a protocol by Klopp *et al.*[12]

**k) NMR measurements of $^{13}C/^{15}N$-labeled RF2**

All NMR measurements were carried out at 30 ˚C using a Bruker 800 MHz NMR spectrometer. The backbone resonance assignments were obtained from the analysis of 3D HNCO, HN(CA)CO, HNCA, HNCACB, together with NOESY-$^{15}N$-HSQC and TOCSY-$^{15}N$-HSQC. Chemical shift assignments were deposited in the BMRB under the accession number 53251 and the title EF2-RF2. Spectra are available on Zenodo (DOI: 10.5281/zenodo.1578834).

**l) Spectral overlap calculations**

The spectral overlap calculations were performed using pre-processed emission and absorption spectra. Spectra were truncated to the range of 280–600 nm. The area under each spectrum was normalized to 1 using trapezoidal integration. Spectral overlap was calculated as the integral of the scalar product between the normalized $Ln^{3+}$ absorption spectrum, and the normalized amino acid emission spectrum, resulting in a dimensionless overlap value between 0 and 1. These calculations were performed in MATLAB.

The absorption spectra of $Eu^{3+}$ and $Tb^{3+}$ were recorded on a Cary 50 UV/Vis spectrophotometer using a 400 µL quartz cuvette. Spectra were acquired in dual beam mode and baseline corrected by blanking with water. Both lanthanide ions were provided as 1 M chloride salt solutions.

Fluorescence emission spectra of the amino acids were recorded in a 400 µL quartz cuvette using a Cary Eclipse fluorescence spectrophotometer. The excitation slit width was set to 10 nm and the emission slit width to 1.5 nm. For all amino acids except mCNP and mCNF, a 295−1100 nm long-pass emission filter was used to suppress scattered excitation light and harmonic doubling. For mCNP and mCNF, the emission filter was left open.

### Supplementary figures

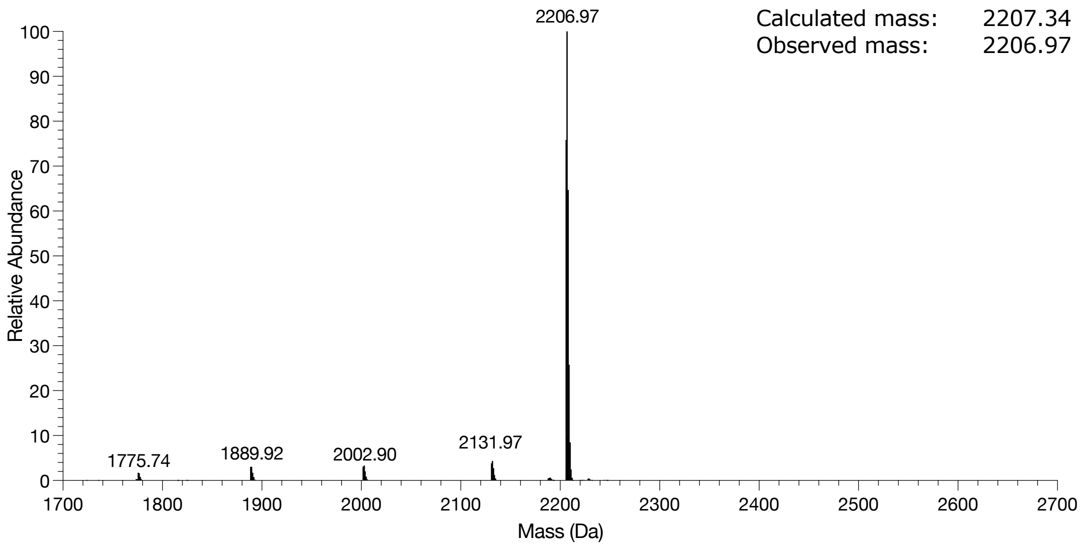


**Figure S1.** Intact protein mass spectrometry of LBT.

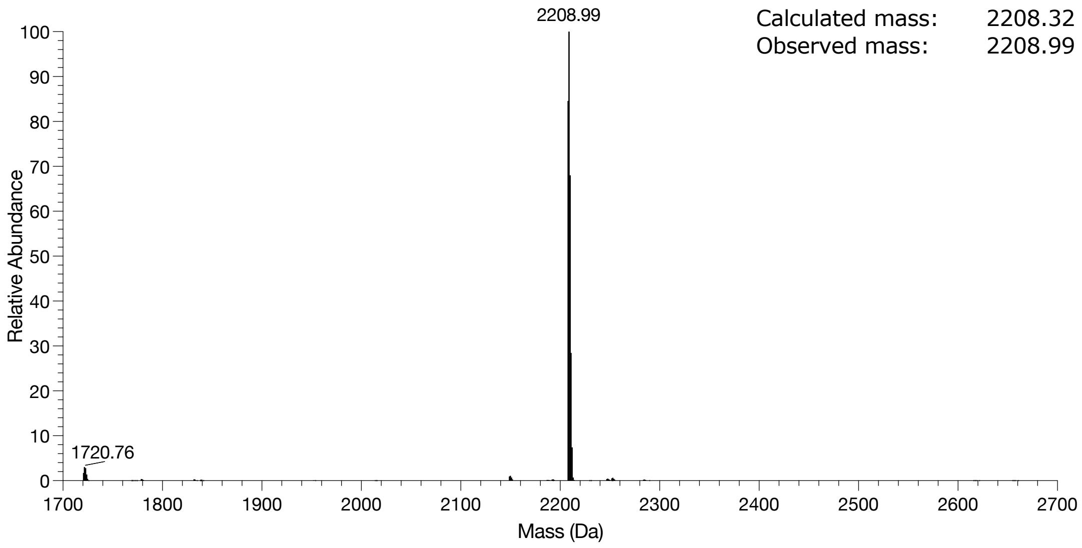


**Figure S2.** Intact protein mass spectrometry of LBT 4AW.

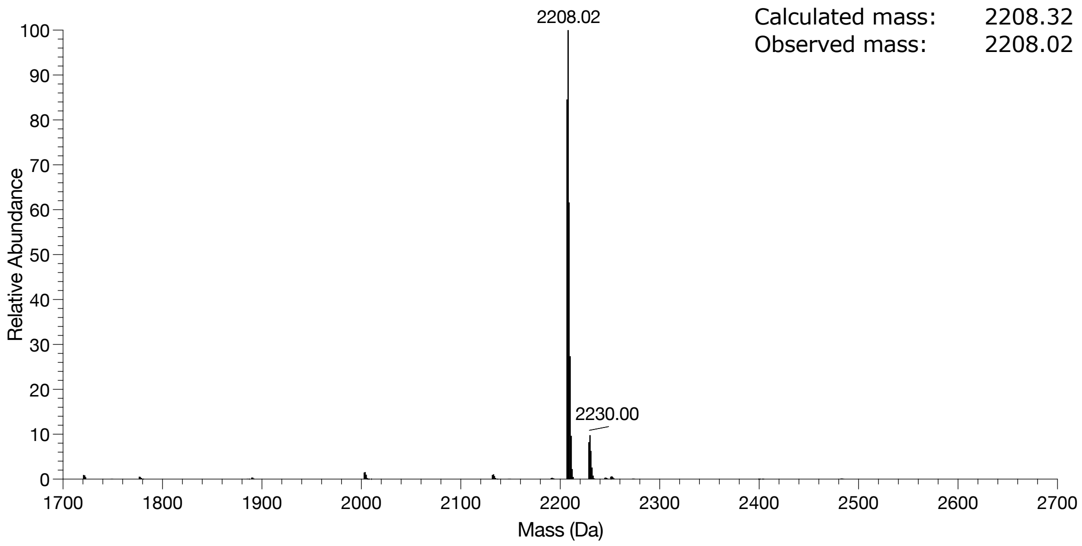


**Figure S3.** Intact protein mass spectrometry of LBT 6AW.

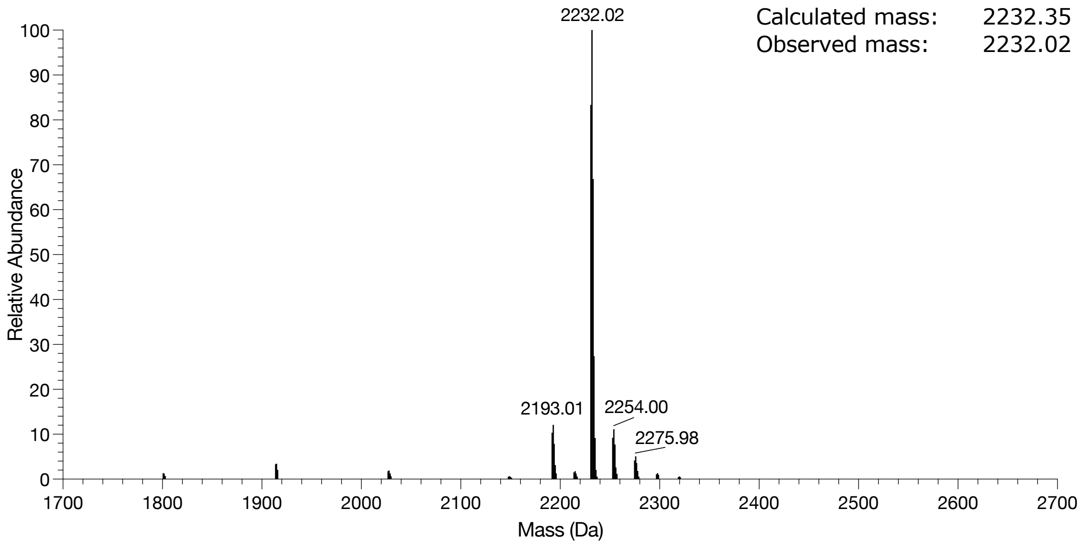


**Figure S4.** Intact protein mass spectrometry of LBT 4CNW.

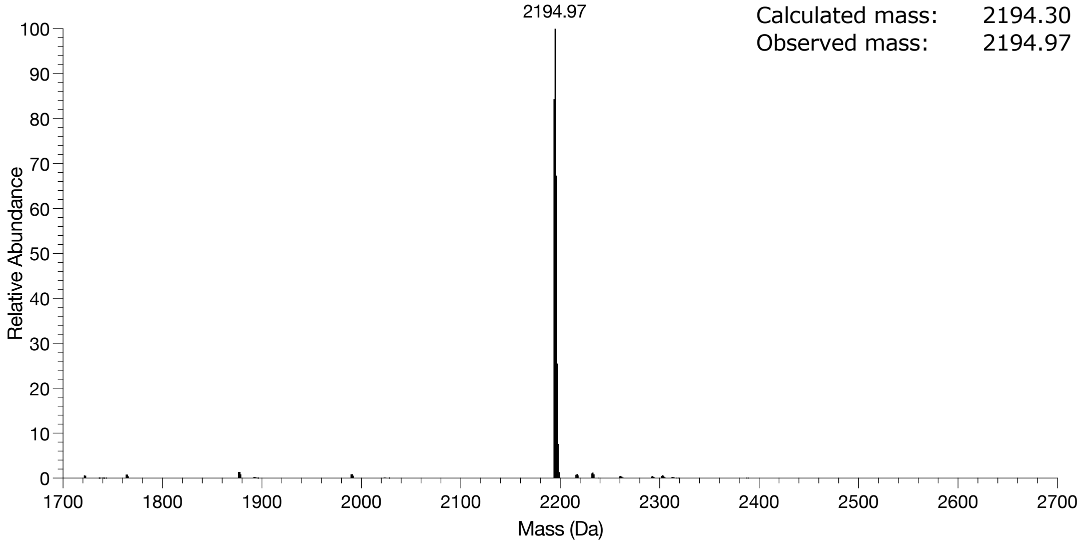


**Figure S5.** Intact protein mass spectrometry of LBT mCNP.

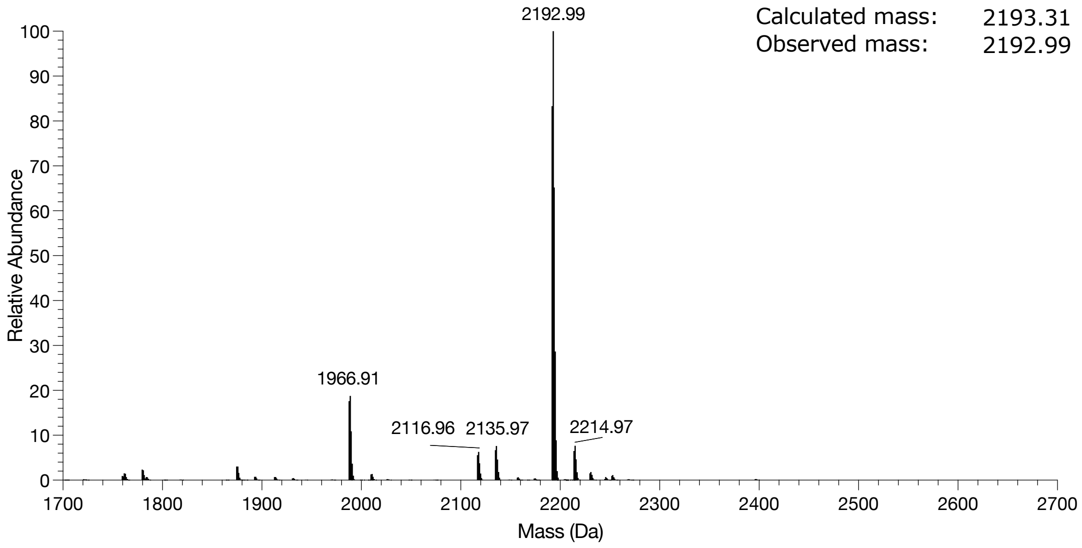


**Figure S6.** Intact protein mass spectrometry of LBT mCNF.

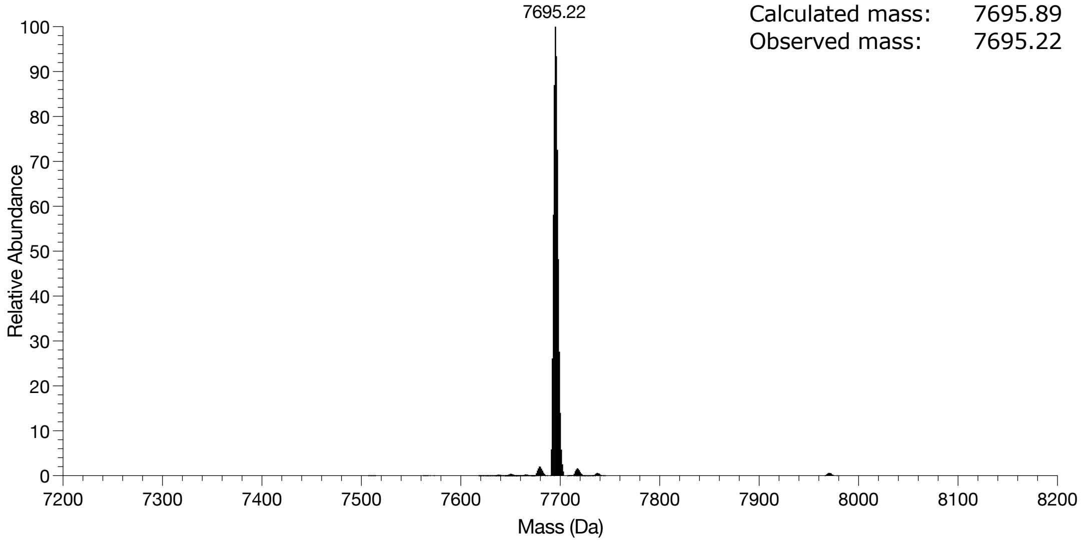


**Figure S7.** Intact protein mass spectrometry of the EF2 variant RF1.

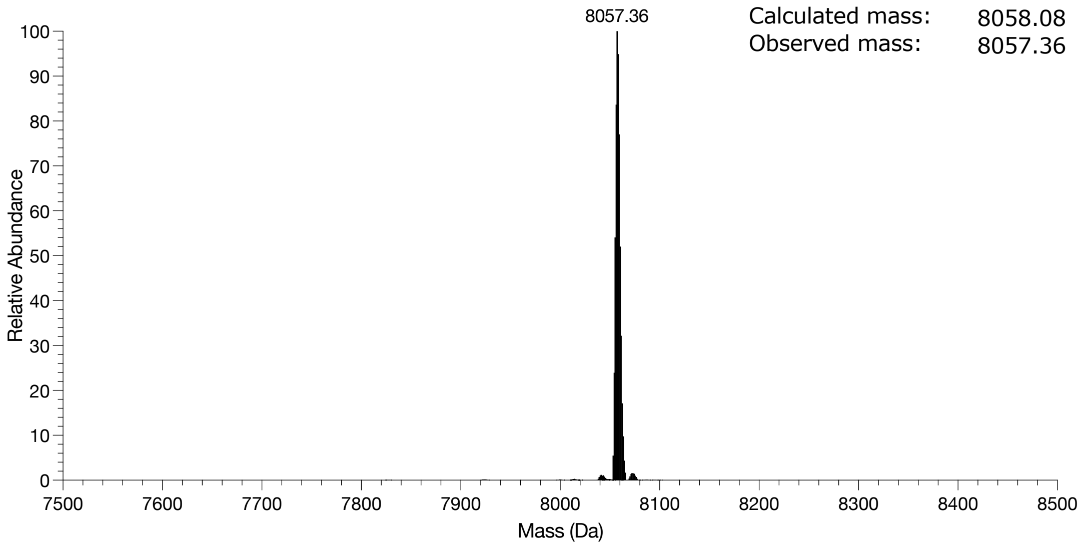


**Figure S8.** Intact protein mass spectrometry of the EF2 variant RF2.

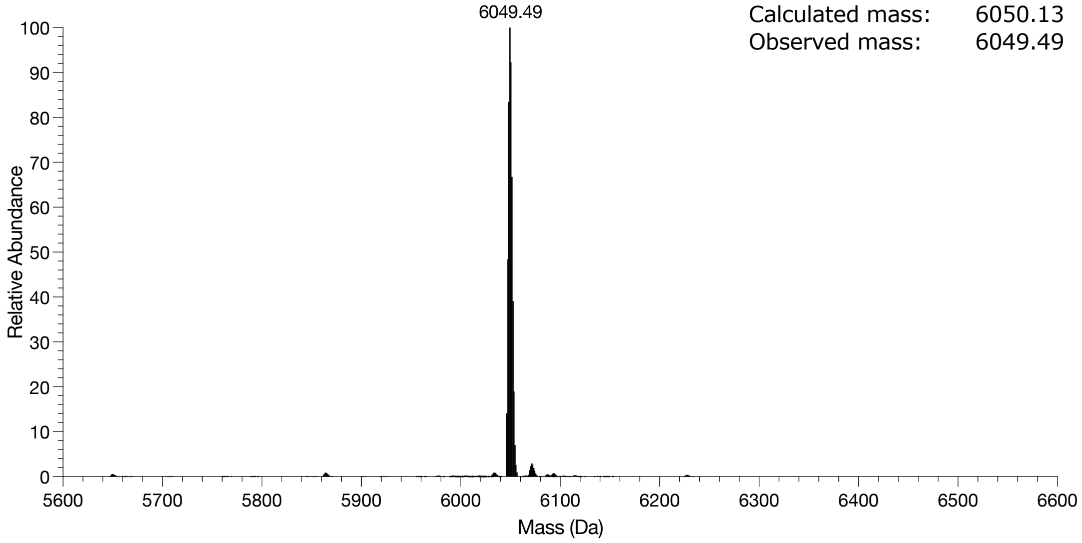


**Figure S9.** Intact protein mass spectrometry of the EF2 variant RF3.

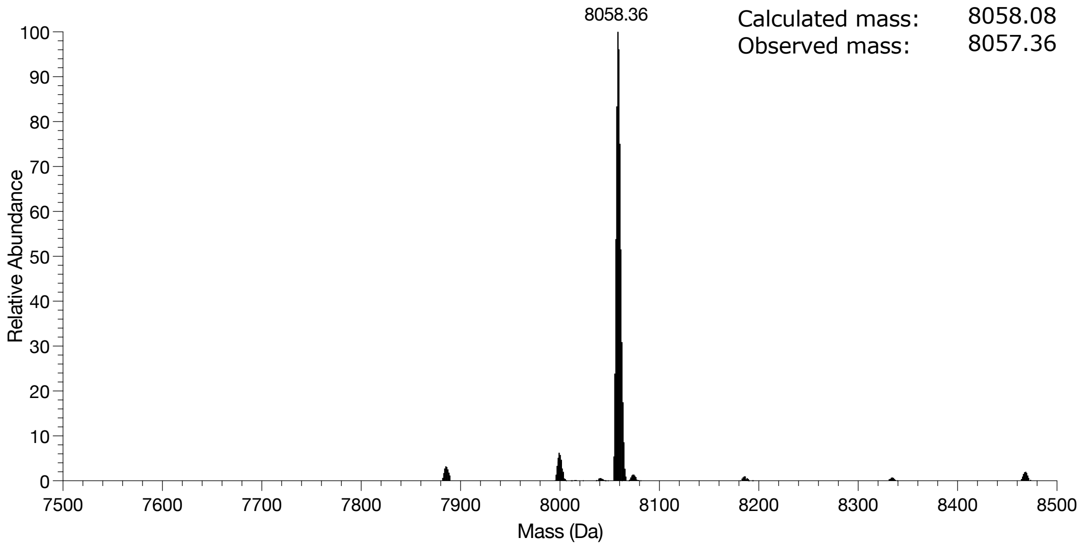


**Figure S10.** Intact protein mass spectrometry of the RF2 42 6AW.

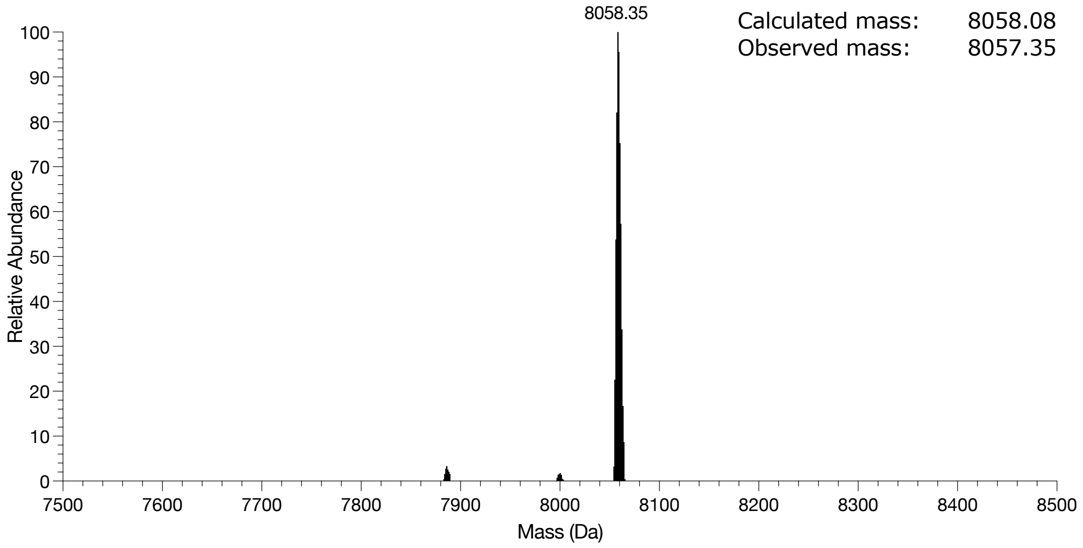


**Figure S11.** Intact protein mass spectrometry of the RF2 50 6AW.

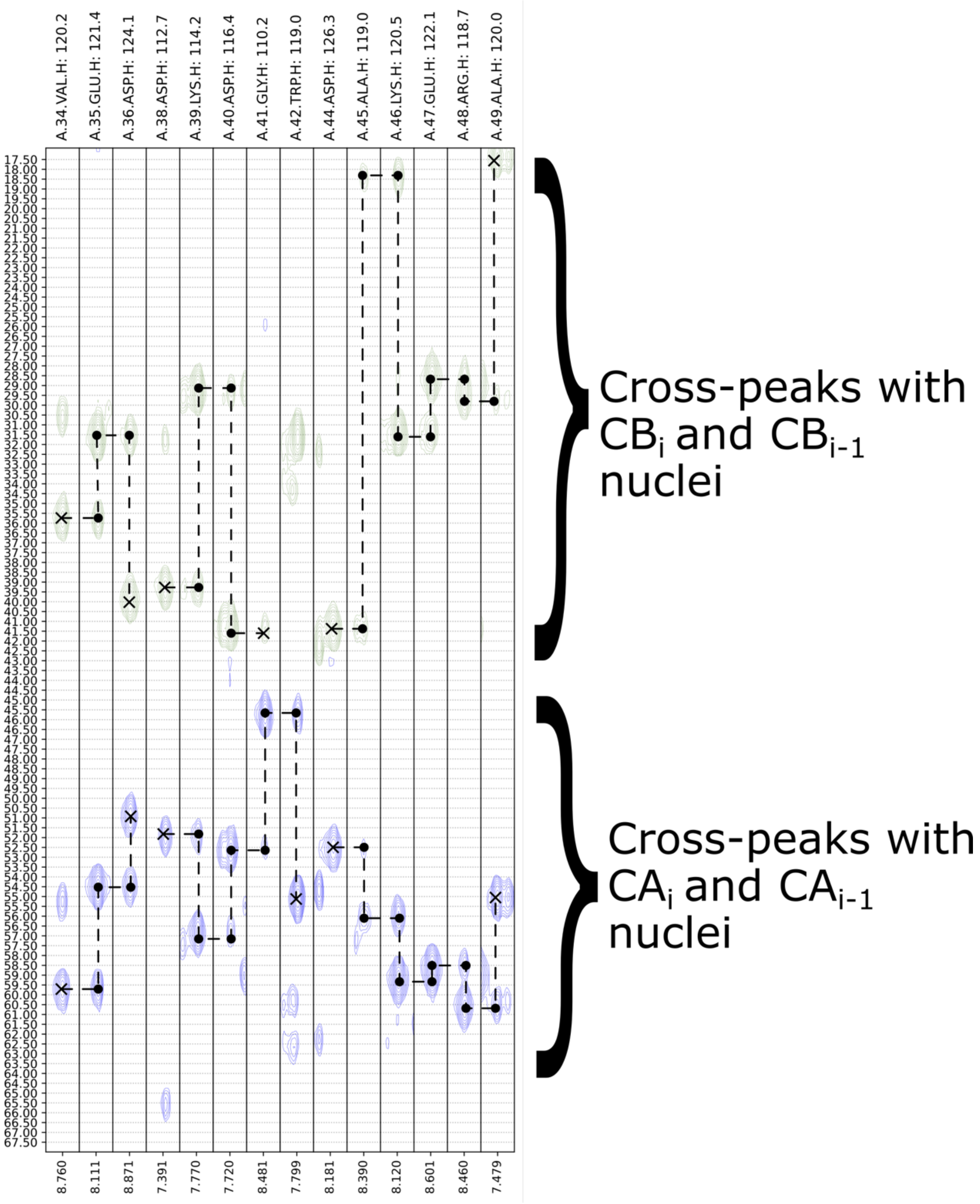


**Figure S12.** HNCACB data of $^{15}N/^{13}C$-labeled RF2. The strip plots are of the 3D HNCACB spectrum of the lanthanide-binding EF-Hand of RF2 including 4 flanking residues (residues 34–49): VEDPDKDGWLDAKERA, where underlined residues do not have assigned NH peaks. The strips illustrate the sequential $CB_i$-$CB_{i-1}$ peaks along the sequence.

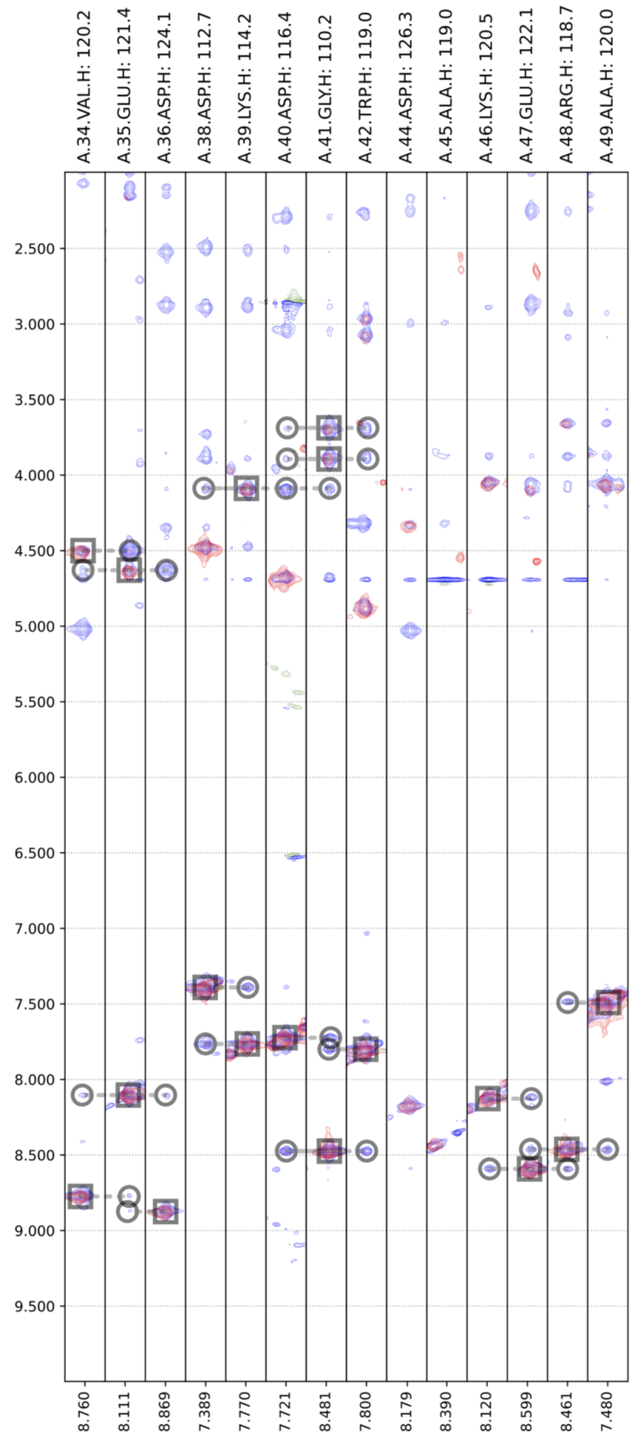


**Figure S13.** Overlay of $^{13}$C-decoupled 3D NOESY-$^{15}$N-HSQC (blue) and TOCSY-$^{15}$N-HSQC (red) spectra of $^{15}$N/$^{13}$C-labeled RF2. The strips belong to the lanthanide-binding EF-Hand of RF2 and 4 flanking residues (res 34–49) VEDPDKDGWLDAKERA, where underlined residues do not have assigned NH peaks. Intra- and inter-residual cross-peaks are identified by squares and circles, respectively.

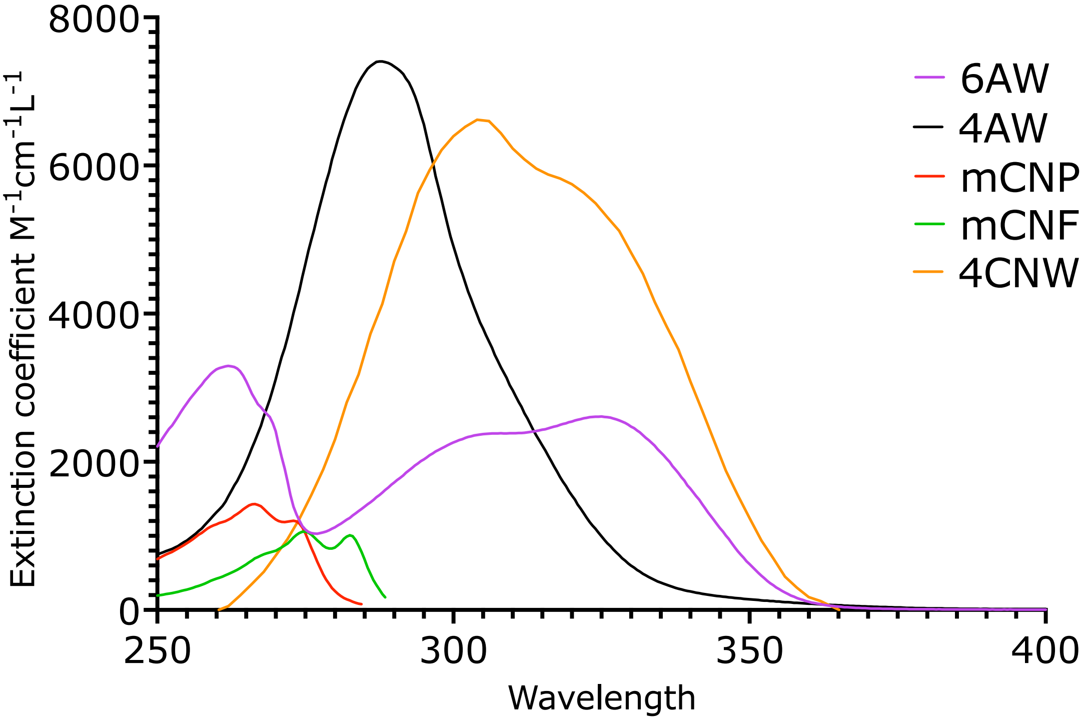


**Figure S14.** Absorbance spectra of the fluorescent ncAAs used in this study.

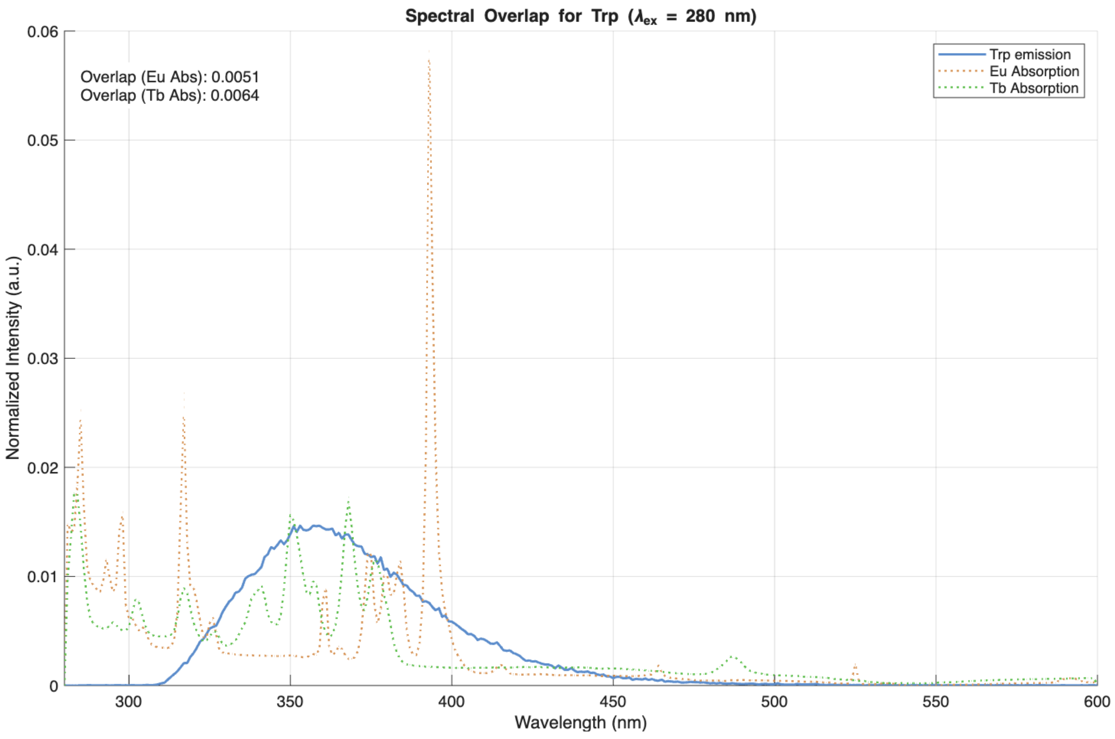


**Figure S15.** Spectral overlap of the Trp emission (free amino acid in Buffer E) with the absorption of $Eu^{3+}$ and $Tb^{3+}$ ($LnCl_3$ in MOPS buffer). Areas of each spectrum are normalized to 1 before plotting.

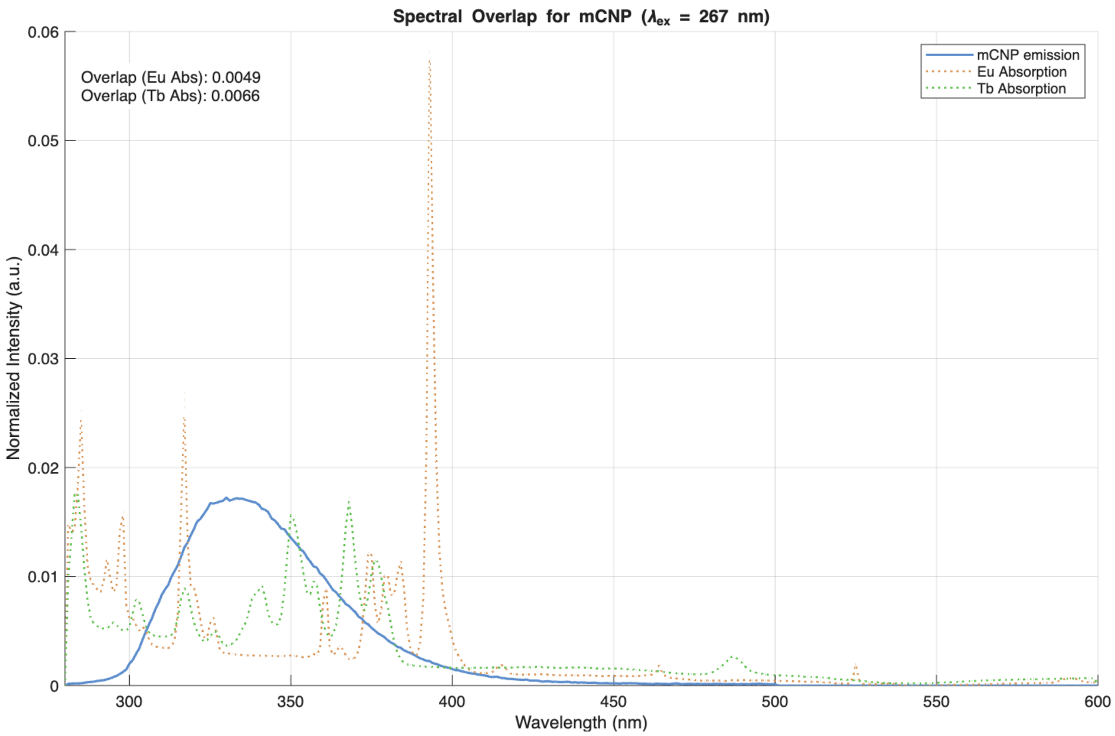


**Figure S16.** Spectral overlap of mCNP emission (free amino acid in Buffer E) with the absorption of $Eu^{3+}$ and $Tb^{3+}$ ($LnCl_3$ in MOPS buffer). Areas of each spectrum are normalized to 1 before plotting.

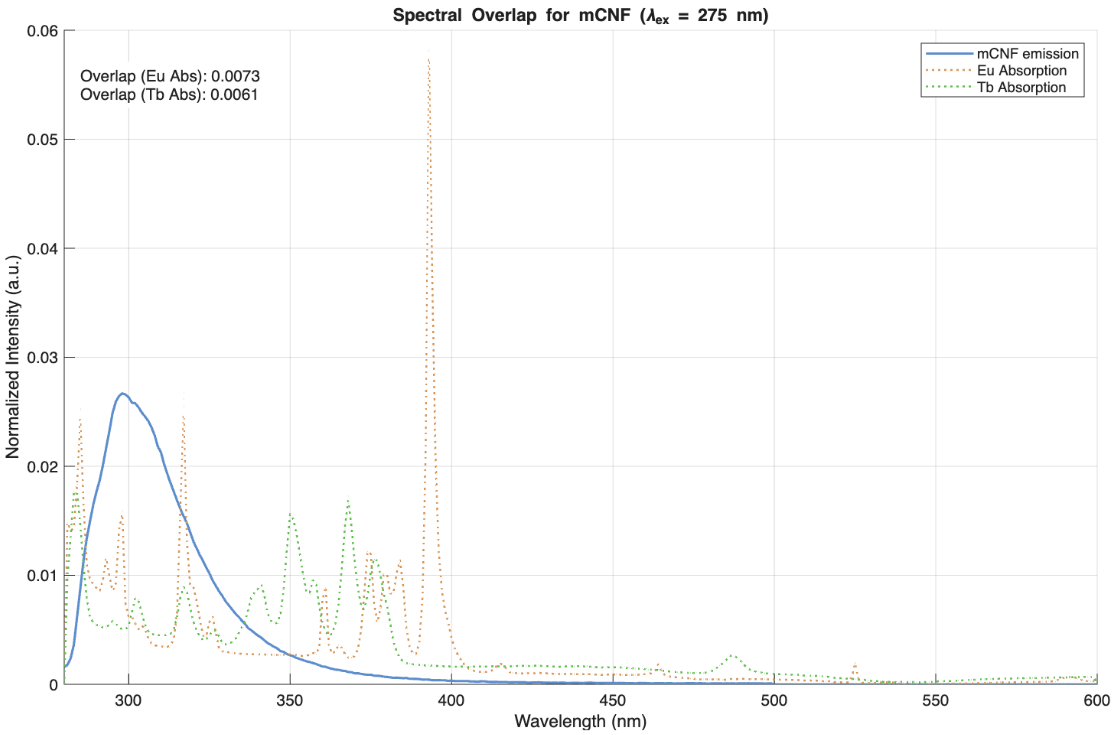


**Figure S17.** Spectral overlap of mCNF emission (free amino acid in Buffer E) with the absorption of $Eu^{3+}$ and $Tb^{3+}$ ($LnCl_3$ in MOPS buffer). Areas of each spectrum are normalized to 1 before plotting.

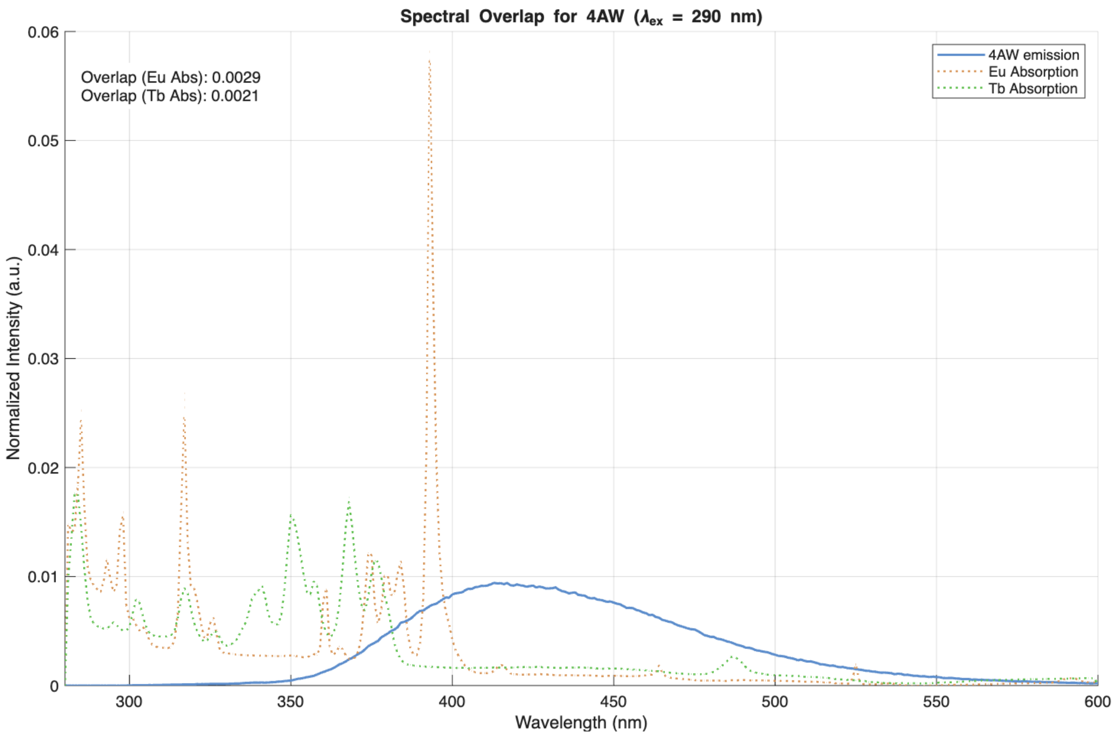


**Figure S18.** Spectral overlap of 4AW emission (free amino acid in Buffer E) with the absorption of $Eu^{3+}$ and $Tb^{3+}$ ($LnCl_3$ in MOPS buffer). Areas of each spectrum are normalized to 1 before plotting.

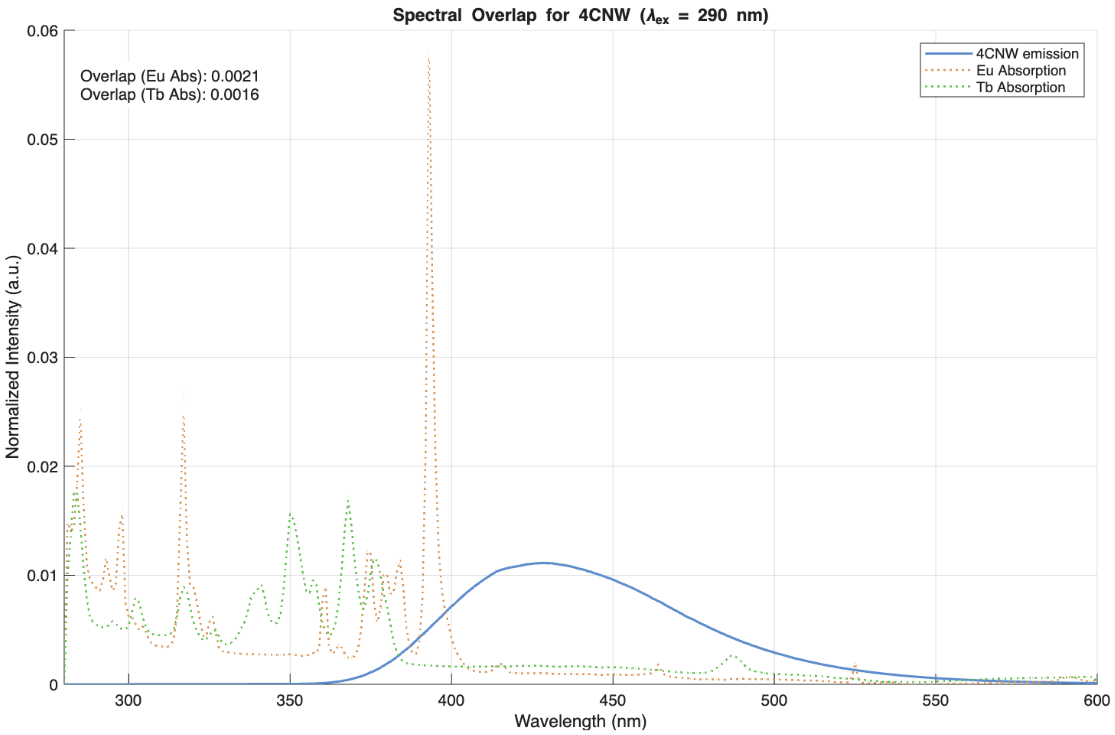


**Figure S19.** Spectral overlap of 4CNW emission (free amino acid in Buffer E) with the absorption of $Eu^{3+}$ and $Tb^{3+}$ ($LnCl_3$ in MOPS buffer). Areas of each spectrum are normalized to 1 before plotting.

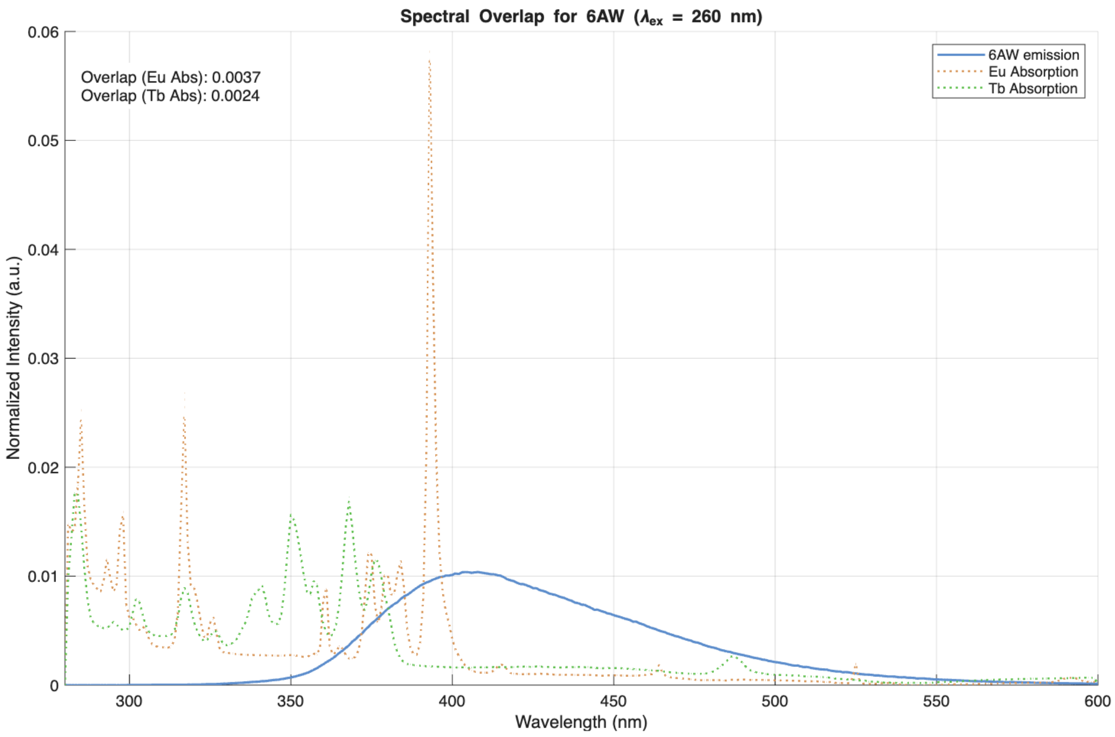


**Figure S20.** Spectral overlap of 6AW emission (free amino acid in Buffer E) with the absorption of $Eu^{3+}$ and $Tb^{3+}$ ($LnCl_3$ in MOPS buffer). Areas of each spectrum are normalized to 1 before plotting.

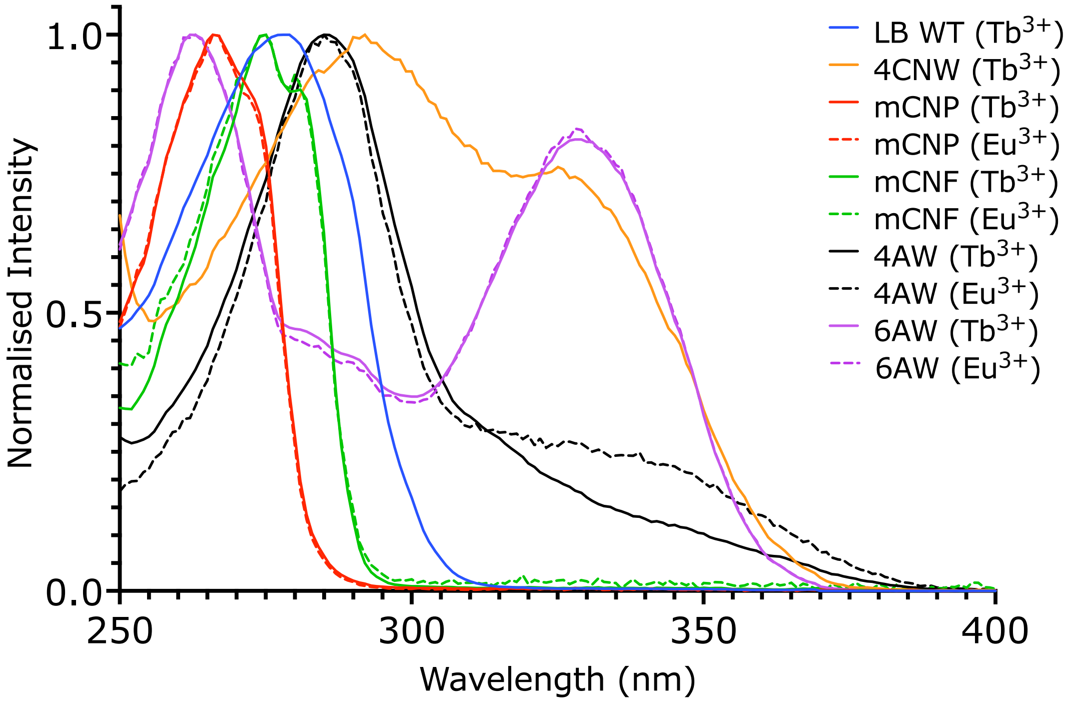

**Figure S21.** Excitation scans of the LBT complexes with $Tb^{3+}$ and $Eu^{3+}$. Emission at 545 nm or 615 nmwas integrated for 5000 µs after a delay of 100 µs, using an excitation slit width of 5 nm and an emission slit width of 10 nm. Protein and lanthanide chloride were at 1 µM and 2 µM, respectively. The samples were prepared in buffer E. Each spectrum is normalized to the maximum intensity.

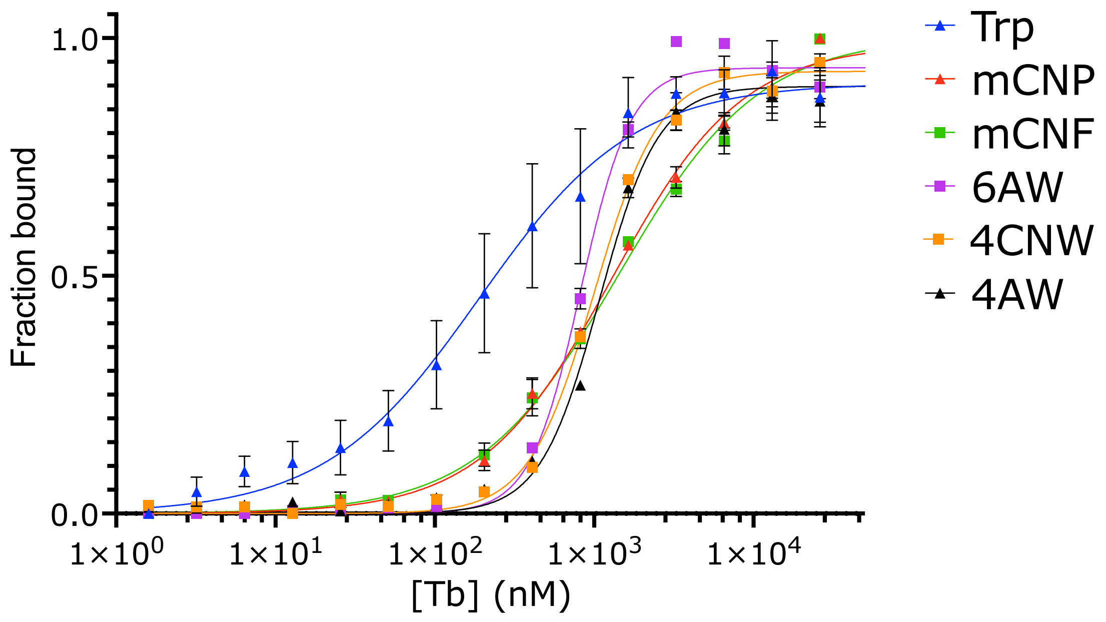


**Figure S22.** Binding affinity determination by titration of LBT variants with $Tb^{3+}$.

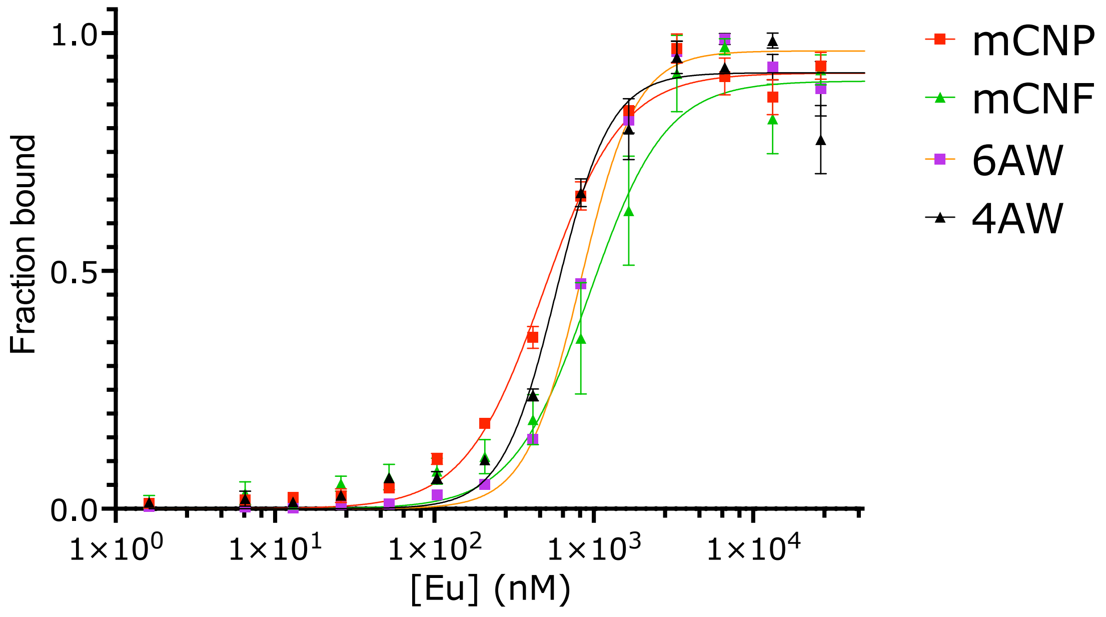

**Figure S23.** Binding affinity determination by titration of LBT variants with $Eu^{3+}$.

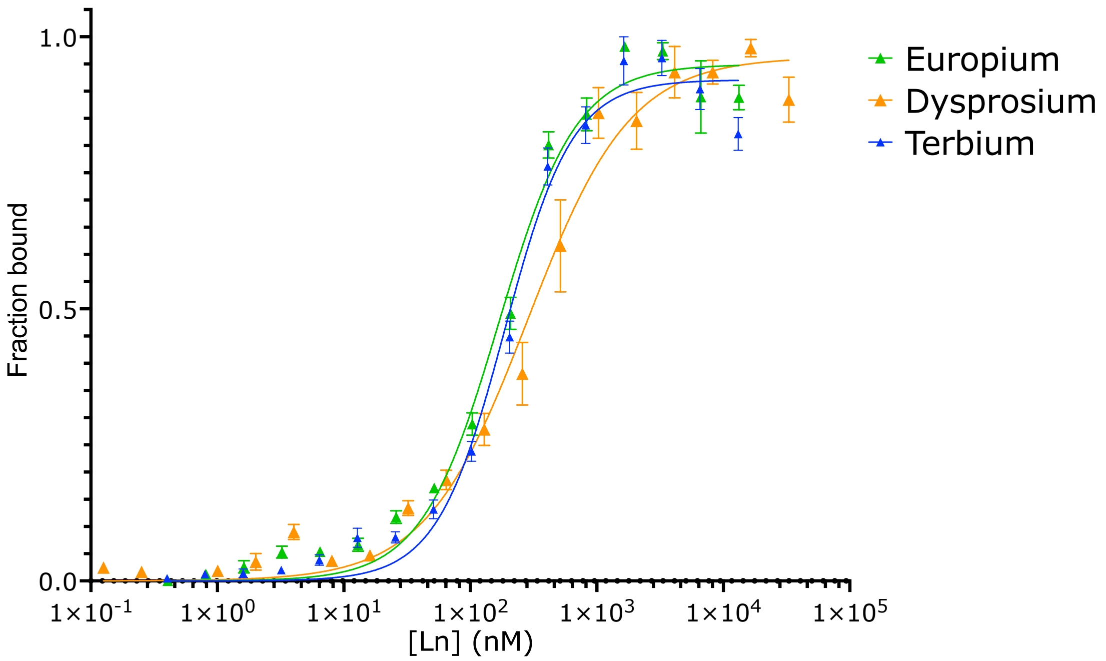


**Figure S24.** Binding affinity determination by titration of RF2 6AW complexes with $Tb^{3+}$, $Eu^{3+}$, and $Dy^{3+}$.

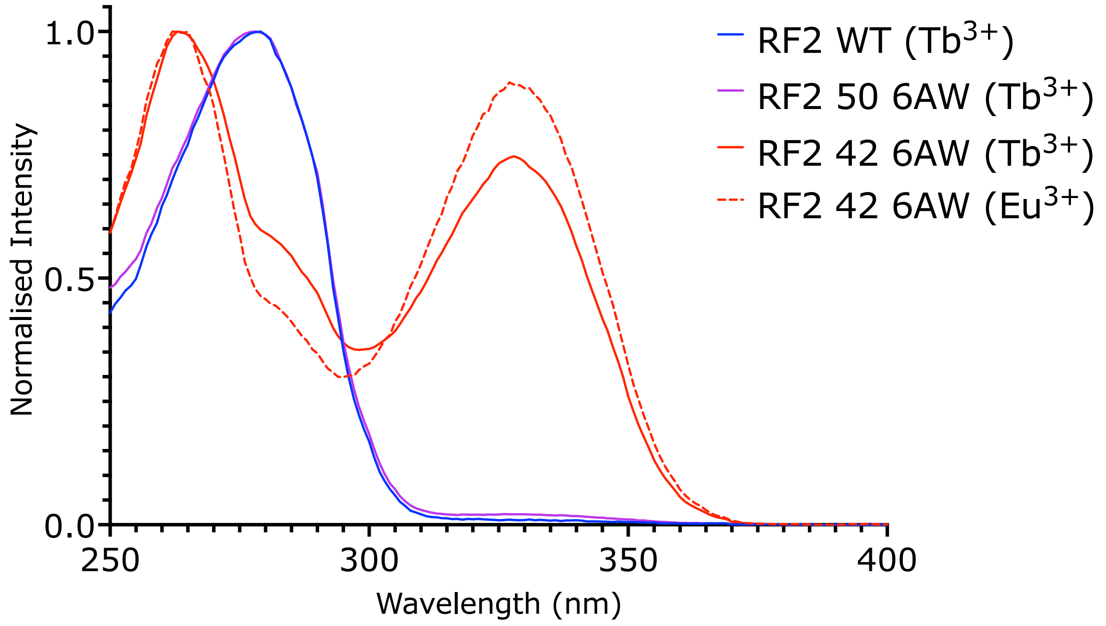


**Figure S25.** Excitation scans of the RF2 complexes with $Tb^{3+}$ and $Eu^{3+}$. Emission at 545 nm or 615 nm was integrated for 5000 µs after a delay of 100 µs, using an excitation slit width of 5 nm and an emission slit width of 10 nm. Protein and lanthanide chloride were at 1 µM and 2 µM, respectively. The samples were prepared in buffer E. Each spectrum is normalized to the maximum intensity.

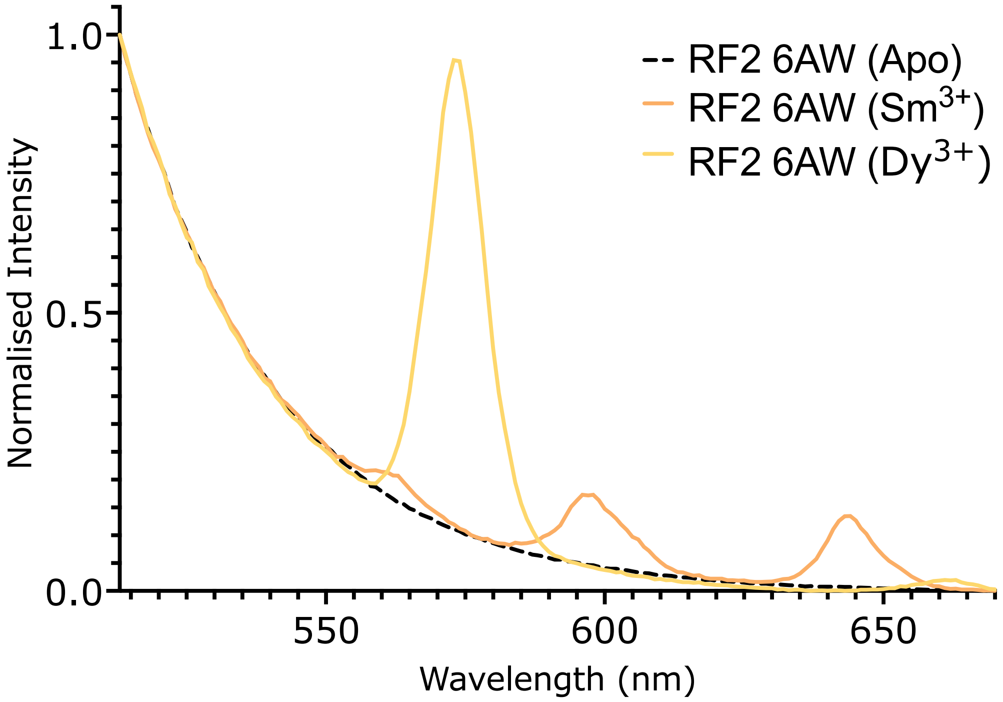


**Figure S26.** Fluorescence emission spectra of RF2 6AW with $Sm^{3+}$ and $Dy^{3+}$. Each spectrum is normalized to the intensity at 520 nm.

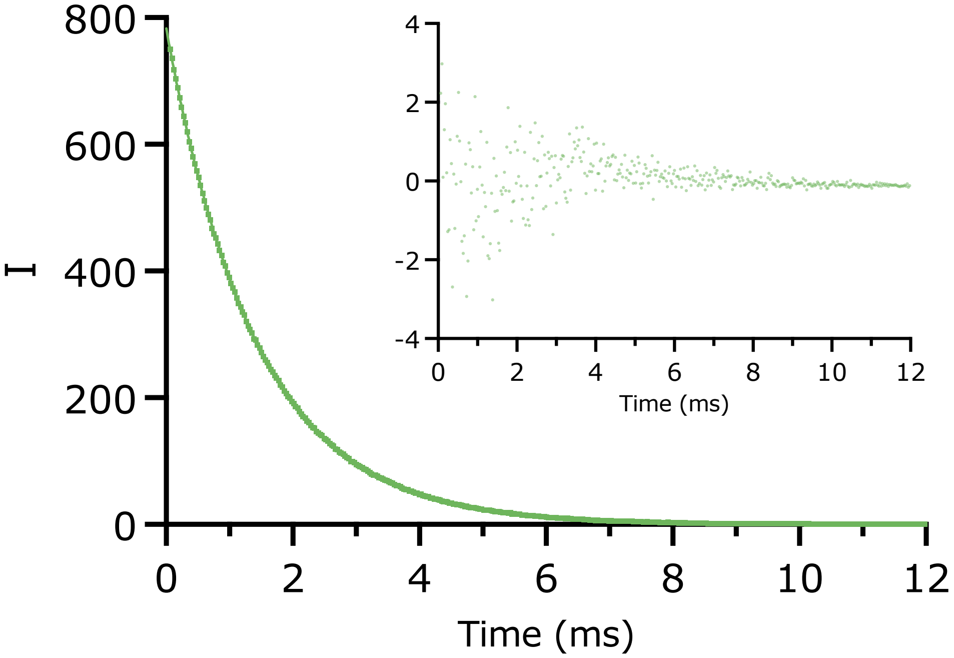

**Figure S27.** Luminescence lifetime measurement of RF2 with $Tb^{3+}$. The excitation wavelength was 280 nm and emission was recorded at 545 nm. Inset: residuals of the experimental values from the curve fit. The best-fit value for $\tau_{Tb}$ is 1.4 ms.

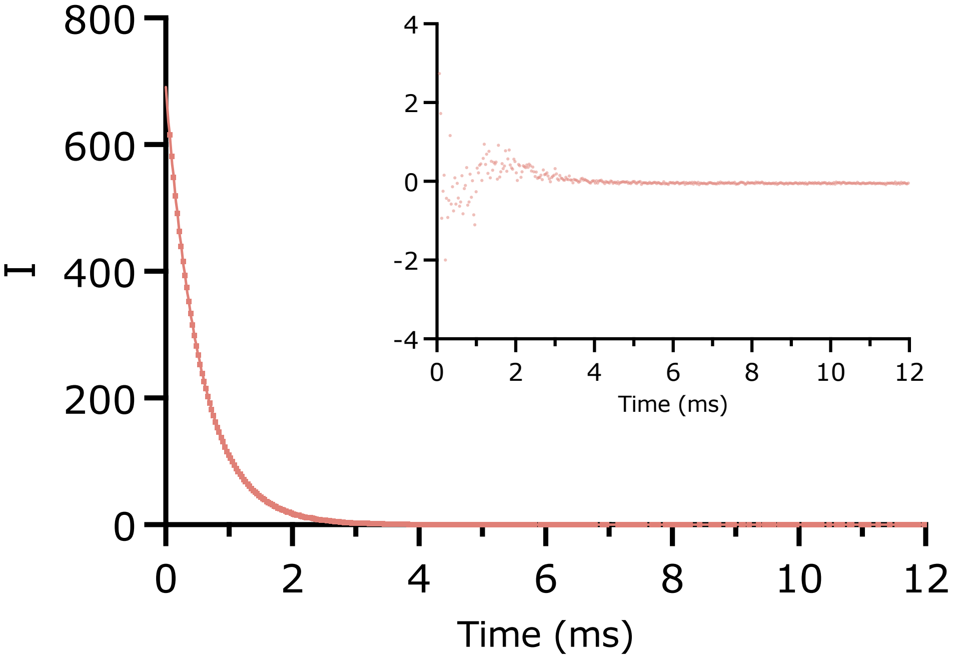


**Figure S28.** Luminescence lifetime measurement of RF2 6AW with $Eu^{3+}$. The excitation wavelength was 330 nm and emission was recorded at 615 nm. Inset: residuals of the experimental values from the curve fit. The best-fit value for $\tau_{Eu}$ is 0.54 ms.

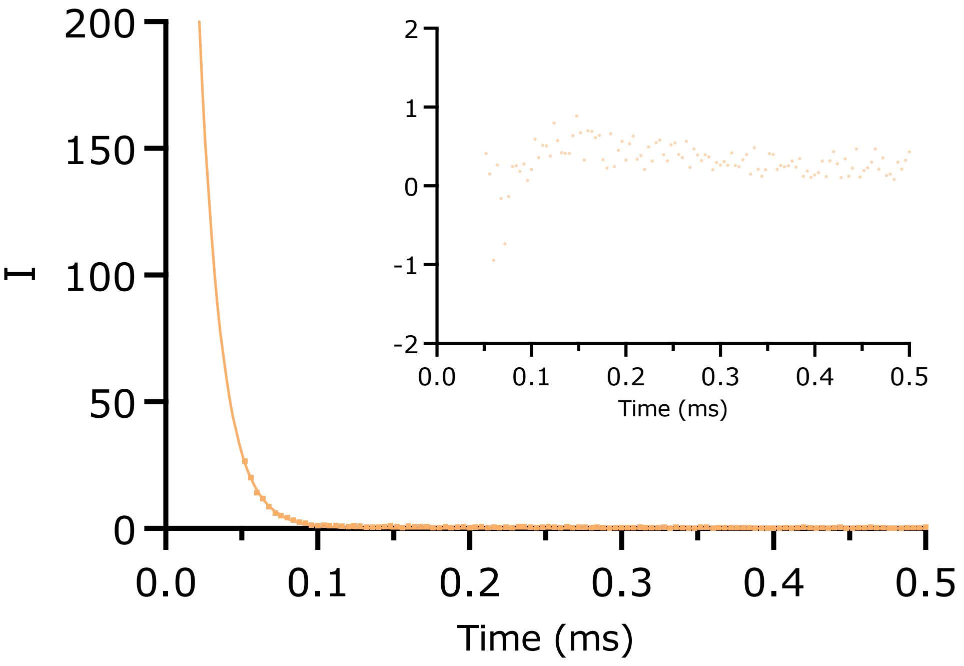


**Figure S29.** Luminescence lifetime measurement of RF2 6AW with $Sm^{3+}$. The excitation wavelength was 265 nm and emission was recorded at 650 nm. The instrument gain was set to 1000. Inset: residuals of the experimental values from the curve fit. The best-fit value for $\tau_{Sm}$ is 12 μs.

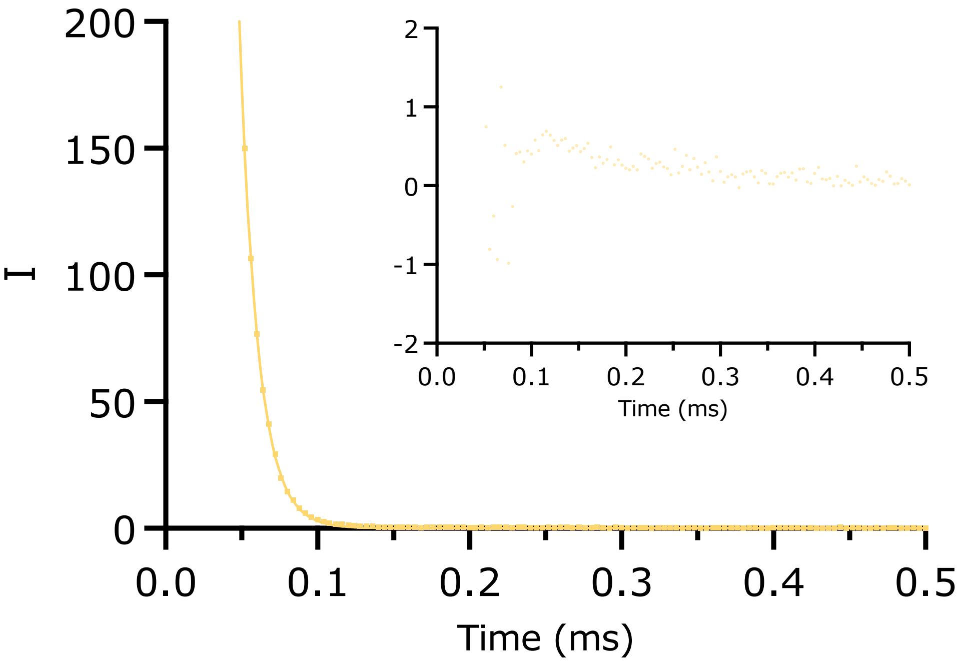

**Figure S30.** Luminescence lifetime measurement of RF2 6AW with $Dy^{3+}$. The excitation wavelength was 265 nm and emission was recorded at 570 nm. Inset: residuals of the experimental values from the curve fit. The best-fit value for $\tau_{Dy}$ is 14 μs.

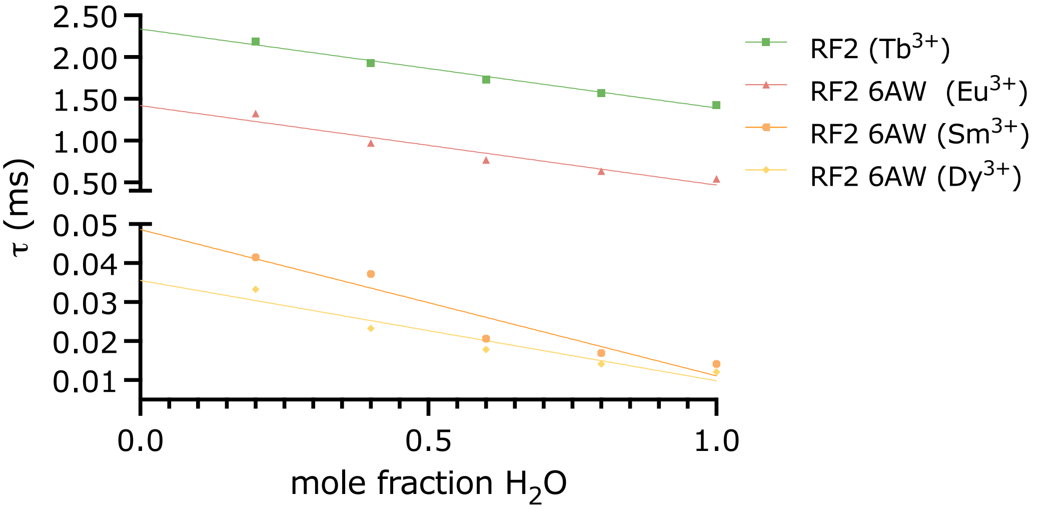


**Figure S31.** $D_2O$ titrations of RF2/$Ln^{3+}$ complexes. The luminescence lifetimes are plotted versus increasing $H_2O$ concentrations. Extrapolation to a $H_2O$ mole fraction of 0 gives the lifetime in $D_2O$.

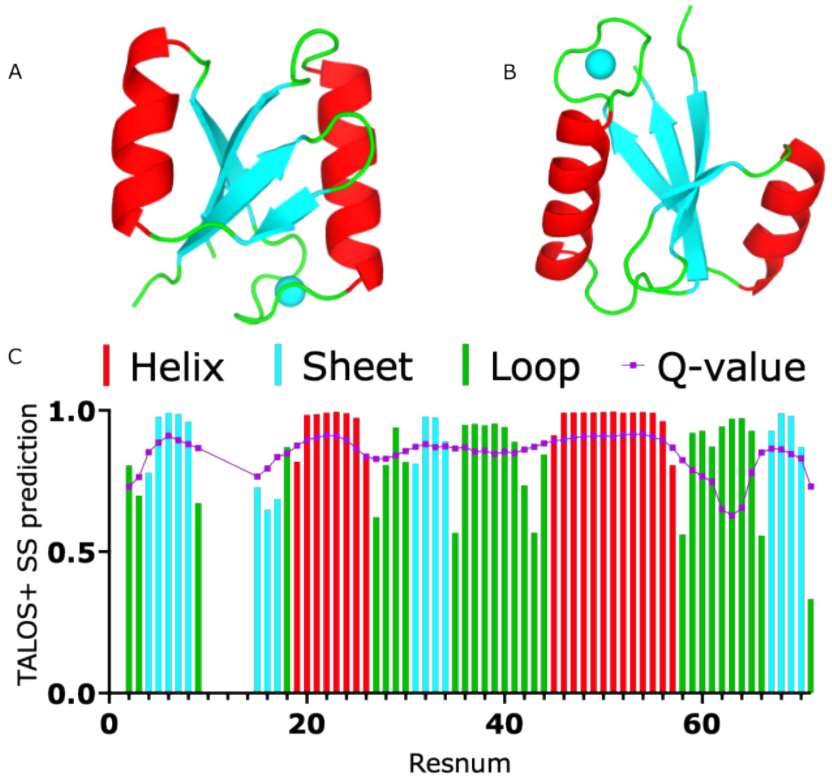

**Figure S32.** Comparison of the secondary structure of the AlphaFold3 model and the prediction made from the assigned chemical shifts. (A) and (B) Two different orientations of the AlphaFold3 model of RF2 generated with a $Ca^{2+}$ ion (cyan). The cartoon representation is colored by secondary structure: red-helix, cyan-sheet, green-loop. (C) Output from the TALOS+ webserver.[13] Colored bars indicate the secondary structure prediction associated with the residue number. The Q-value (purple) indicates the confidence for that prediction.

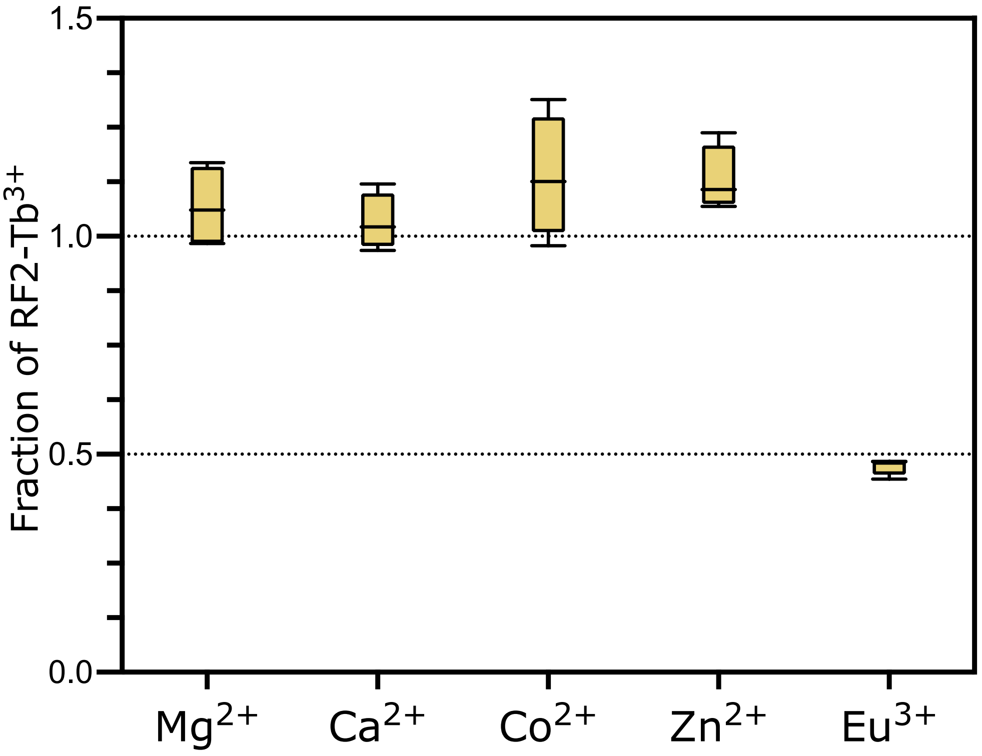


**Figure S33.** Competition experiment with $M^{2+}$ ions. Fold-change in the luminescence intensity (gold) of RF2 (1 µM) samples with $Tb^{3+}$ (2 µM) and competitor (2 µM) in 20 mM MOPS pH 7.0, 50 mM KCl. Each data point represents the ratio of integrated emission intensity (collected at 545 nm emission with a 100 µs delay and 1 ms integration window, following excitation at 280 nm) normalized to the intensity of a sample containing only $Tb^{3+}$. Boxes indicate the mean and range of four technical replicates.

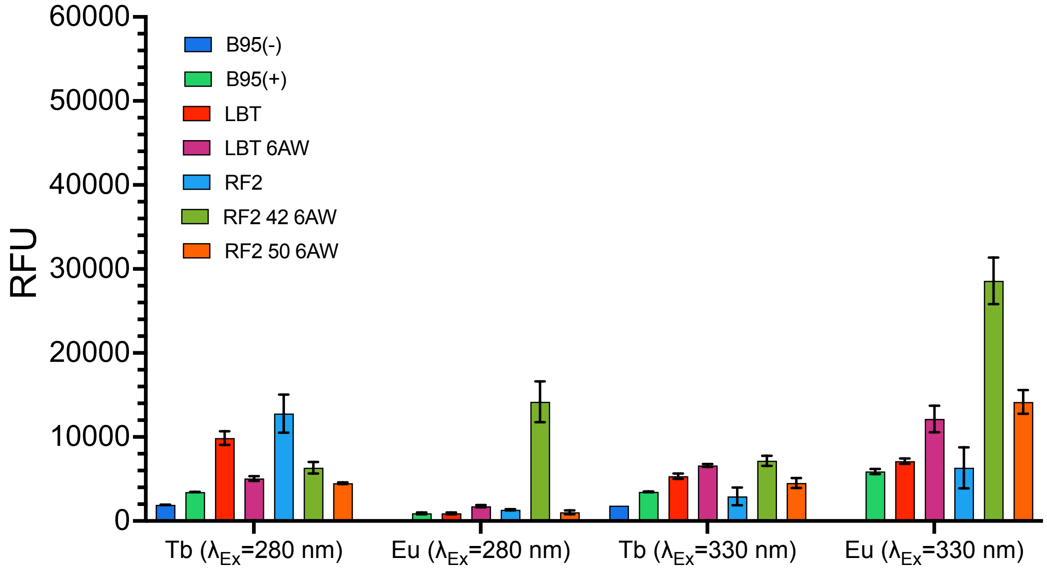


**Figure S34.** *In vivo* sensitization of lanthanide luminescence. Untransformed B95 cells with and without 6AW (B95(+), B95(-)), transformed B95 cells with a WT LBT or RF2 plasmid, or the two-plasmid expression system for installing 6AW were grown to $OD_{600}$ = 0.6–0.8 in LB media and supplemented with 1 mM IPTG. Cultures with cells containing the G1(6AW)RS plasmid, as well as the B95(+) cells were supplemented with 1 mM 6AW. Cells were cultured overnight at RT and resuspended to $OD_{600}$ = 1.0 in 150 mM NaCl. The standardized cell resuspensions were then resuspended again in 150 mM NaCl containing 25 mM $EuCl_3$ or 25 mM $TbCl_3$. Readings are integrated emission intensities over 1000 µs following a 600 µs delay. Emission intensities were $\lambda_{Em}$ = 545 nm for Tb, and $\lambda_{Em}$ = 615 nm for Eu. Datasets for Eu- and Tb-incubated cells were recorded with different gain settings for the detector. Therefore the values between them are not directly comparable.

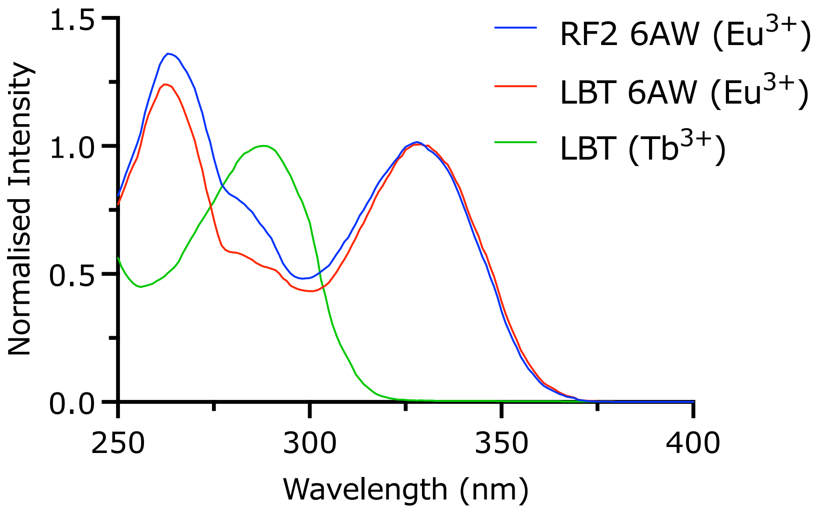

**Figure S35.** Comparison of excitation spectra for RF2 6AW/$Eu^{3+}$, LBT 6AW/$Eu^{3+}$ and LBT/$Tb^{3+}$. RF2 6AW/$Eu^{3+}$ (blue) and LBT 6AW/$Eu^{3+}$ have been normalized to the intensity of 330 nm, where the contribution of canonical amino acids to sensitization is negligible. LBT/$Tb^{3+}$ has been normalized to the intensity at 280 nm, as in Figure S21.

### Supplementary tables

**Table S1.** Extinction coefficients of the amino acids used in this study

| Amino acid | $\lambda_{ex}$ (nm) | Extension coefficient[a] ($M^{-1}$ $cm^{-1}$) |
|---|---|---|
| 4-CN-Trp[a,b] | 290 | 4708 ± 58 (at 290 nm)<br>2304 ± 48 (at 280 nm) |
| 6-Aza-Trp | 265 | 3087 ± 8 (at 265 nm)<br>1013 ± 4 (at 280 nm) |
| 4-Aza-Trp | 290 | 7335 ± 4 (at 290 nm)<br>6272 ± 4 (at 280 nm) |
| *m*-cyano-pyridylalanine | 265 | 1386 ± 9 (at 265 nm)<br>255 ± 9 (at 280 nm) |
| *m*-cyano-phenylalanine | 275 | 1059 ± 30 (at 275 nm)<br>843 ± 19 (at 280 nm) |

[a] Extinction coefficients at 280 nm are useful for determining protein concentrations based on their amino acid sequence. For comparison, the $\varepsilon_{280}$ value of tryptophan is 5422 $M^{-1}$ $cm^{-1}$.
[b] Value from Abdelkader *et al.*[14]

**Table S2.** Variables used in integrated lifetime calculations[a]

| Protein | $Ln^{3+}$ | $Y_0$ | | Plateau | | *K* | | Integral | | Error (%) |
|---|---|---|---|---|---|---|---|---|---|---|
| LBT | Tb | 259.17 | ± 0.20 | 0.00 | ± 0.07 | 0.38 | ± 0.00 | 682.88 | ± 1.14 | 0.17 |
| LBT | Eu | 3.32 | ± 0.03 | 0.00 | ± 0.00 | 0.71 | ± 0.00 | 4.65 | ± 0.04 | 0.79 |
| LBT mCNP | Tb | 233.43 | ± 0.20 | -0.15 | ± 0.07 | 0.37 | ± 0.00 | 630.27 | ± 1.17 | 0.19 |
| LBT mCNP | Eu | 253.05 | ± 0.18 | -0.01 | ± 0.03 | 0.70 | ± 0.00 | 361.21 | ± 0.48 | 0.13 |
| LBT mCNF | Tb | 156.29 | ± 0.15 | -0.14 | ± 0.05 | 0.37 | ± 0.00 | 423.25 | ± 0.85 | 0.20 |
| LBT mCNF | Eu | 35.33 | ± 0.06 | 0.00 | ± 0.01 | 0.70 | ± 0.00 | 50.39 | ± 0.16 | 0.32 |
| LBT 4AW | Tb | 468.61 | ± 0.92 | 2.16 | ± 0.18 | 0.64 | ± 0.00 | 728.16 | ± 2.67 | 0.37 |
| LBT 4AW | Eu | 61.71 | ± 0.09 | 0.01 | ± 0.02 | 0.72 | ± 0.00 | 85.77 | ± 0.24 | 0.28 |
| LBT 6AW | Tb | 402.22 | ± 0.30 | -0.14 | ± 0.10 | 0.38 | ± 0.00 | 1072.23 | ± 1.71 | 0.16 |
| LBT 6AW | Eu | 268.10 | ± 0.24 | 0.01 | ± 0.04 | 0.71 | ± 0.00 | 377.86 | ± 0.62 | 0.16 |
| LBT 4CNW | Tb | 209.92 | ± 0.31 | 0.16 | ± 0.06 | 0.65 | ± 0.00 | 323.19 | ± 0.88 | 0.27 |
| LBT 4CNW | Eu | 3.77 | ± 0.02 | -0.00 | ± 0.00 | 0.70 | ± 0.01 | 5.38 | ± 0.06 | 1.20 |
| RF1 | Tb | 867.20 | ± 0.48 | 0.17 | ± 0.09 | 0.72 | ± 0.00 | 1207.33 | ± 1.24 | 0.10 |
| RF2 | Tb | 1236.34 | ± 0.51 | 0.02 | ± 0.07 | 0.69 | ± 0.00 | 1783.83 | ± 1.20 | 0.07 |
| RF2 | Eu | 0.67 | ± 0.02 | -0.01 | ± 0.00 | 1.11 | ± 0.04 | 0.61 | ± 0.03 | 4.77 |
| RF3 | Tb | 357.29 | ± 0.59 | 0.28 | ± 0.10 | 0.77 | ± 0.00 | 462.69 | ± 1.38 | 0.30 |
| RF2 42 6AW[b] | Tb | 1027.07 | ± 0.54 | 0.07 | ± 0.10 | 0.68 | ± 0.00 | 1507.04 | ± 1.46 | 0.10 |
| RF2 42 6AW[b] | Eu | 920.91 | ± 0.47 | 0.03 | ± 0.04 | 1.82 | ± 0.00 | 506.04 | ± 0.43 | 0.09 |
| RF2 42 6AW[b] | Dy | 1631.29 | ± 119.71 | 0.02 | ± 0.00 | 56.77 | ± 0.98 | 28.73 | ± 2.17 | 7.54 |
| RF2 50 6AW[b] | Tb | 1200.19 | ± 0.67 | 0.03 | ± 0.10 | 0.71 | ± 0.00 | 1700.41 | ± 1.60 | 0.09 |
| RF2 50 6AW[b] | Eu | 24.18 | ± 0.07 | -0.02 | ± 0.01 | 1.88 | ± 0.01 | 12.84 | ± 0.06 | 0.48 |

[a] Uncertainties were derived from the deviation of the best-fit equation to the data. Final values are rounded to whole numbers in the main text.

[b] The notations used here are different from the main text to facilitate discrimination of the different 6AW positions.

**Table S3.** Amino acid sequences of the proteins used in this study

| Protein | Amino acid sequence[a] |
|---|---|
| NT-LBT-Trp | MASMTGHHHHHHSHTTPWTNPGLAENFMNSFMQGLSSMPGFTASQLDK MSTIAQSMVQSIQSLAAQGRTSPNDLQALNMAFASSMAEIAASEEGGGSLS TKTSSIASAMSNAFLQTTGVVNQPFINEITQLVSMFAQAGMNDVSAGNSEN LYFQGFIDTNNDGWIEGDELLLEG |
| NT-LBT-Amb | MASMTGHHHHHHSHTTPWTNPGLAENFMNSFMQGLSSMPGFTASQLDK MSTIAQSMVQSIQSLAAQGRTSPNDLQALNMAFASSMAEIAASEEGGGSLS TKTSSIASAMSNAFLQTTGVVNQPFINEITQLVSMFAQAGMNDVSAGNSEN LYFQGFIDTNNDGZIEGDELLLEG |
| EF2 RF1-Trp | MASMTGHHHHHHGGSENLYFQGKKVITISEDTPESFEVKMGKGLLKVTVP GKTFIVEDPDKDGWLDAKEIKALLAAAKAATGAKTIEIEDIPK |
| EF2 RF2-Trp | MASMTGHHHHHHGGSENLYFQGRRAYLLRVDPDTLVEISPEEAERLAKTR PVLEVEDPDKDGWLDAKERAWILEHLRETHPDASGAVIVVVD |
| EF2 RF3-Trp | MASMTGHHHHHHGGSENLYFQGKTITTYKFATKEDEKKAKELDPDKDG WLDAKELKKLKKAGLKITKVKTKTIK |
| EF2 RF2-42Z | MASMTGHHHHHHGGSENLYFQGRRAYLLRVDPDTLVEISPEEAERLAKTR PVLEVEDPDKDGZLDAKERAWILEHLRETHPDASGAVIVVVD |
| EF2 RF2-50Z | MASMTGHHHHHHGGSENLYFQGRRAYLLRVDPDTLVEISPEEAERLAKTR PVLEVEDPDKDGWLDAKERAZILEHLRETHPDASGAVIVVVD |

[a] Z indicates the positions of the amber sites.

**Table S4.** Yields of different proteins co-expressed with G1(ncAA)RS in LB media with 1 mM ncAA[a]

| **Amber site** | **G1(ncAA)RS** | **Yield (mg per 1 L cell culture)** |
|---|---|---|
| NT-LBT | - | 31[b] |
| NT-LBT 4AW | 4AWRS[15] | 16[c] |
| NT-LBT 6AW | 6AWRS[15] | 26[c] |
| NT-LBT 4CNW | 4CNWRS[14] | 20[c] |
| NT-LBT mCNP | mCNPRS[16] | 10[c] |
| NT-LBT mCNF | mCNPRS[16] | 15[c] |
| EF2 RF1 | - | 50[b] |
| EF2 RF2 | - | >400[b] |
| EF2 RF3 | - | 10[b] |
| EF2 RF2 42 6AW | 6AWRS | 51[d] |
| EF2 RF2 50 6AW | 6AWRS | 19[e] |

[a] All proteins were expressed and purified as described in the methods section. Yields were calculated by measuring the protein concentration at a known volume and adjusting for expression volume. [b] From 1 L expression volume. [c] From 100 mL expression volume. [d] From 50 mL expression volume. [e] From 30 mL expression volume.

**Table S5.** EF2-RF variants with $Tb^{3+}$

| Protein | Intensity Tb* | Apparent $K_D$ ($Tb^{3+}$) (nM) | Lifetime ($\tau_{Tb}$) |
|---|---|---|---|
| EF2 RF1 | 1207 | 320 ± 50 | 1.4 |
| EF2 RF2 | 1734 | 20 ± 3 | 1.4 |
| EF2 RF3 | 462 | 3300 ± 130 | 1.3 |

**Table S6.** Determination of q-values from $D_2O$ titrations[a]

| Complex | $\tau_{H_2O}$ | $\tau_{D_2O}$ | $q$ | Uncertainty on $q$ |
|---|---|---|---|---|
| RF2 WT Tb | 1.42 ms | 2.34 ms | 1.07 | ±0.3 |
| RF2 42 6AW Eu | 0.54 ms | 1.42 ms | 1.06 | ±0.3 |
| RF2 42 6AW Sm | 12.12 µs | 48.64 µs | 1.30 | ±0.5 |
| RF2 42 6AW Dy | 14.18 µs | 35.63 µs | 1.31 | ±0.5 |

[a] The q values were determined using the modified Horrocks equation as described above. Uncertainties are from errors in the empirical parameters used to determine the q value.
